# Fruitless produces diverse neuronal sex differences by editing lineage-programmed chromatin landscapes

**DOI:** 10.64898/2026.04.14.718444

**Authors:** Margarita V. Brovkina, E. Josephine Clowney

## Abstract

A major question in metazoan development is how a limited set of DNA-binding transcription factors generate the much larger number of cell fates and states in the organism. *In vivo*, individual transcription factors are joined by varied sets of collaborating or competing proteins across cellular contexts. Brains provide particularly diverse transcriptional milieus across thousands of neuron types, where each cell is a phenotypic readout of intrinsic transcriptional regulation over developmental time. Today’s methods allow interrogation of individual cells, transforming the diversity of neurons in the brain from an experimental hindrance to a benefit in interrogation of combinatorial transcription factor action. Here, we use the BTB domain transcription factor Fruitless as a model to determine how a single factor produces cell-type specific regulatory effects, in this case sexual dimorphisms. In *Drosophila melanogaster*, Fruitless is transcribed in a precise neuronal repertoire (irrespective of sex) but translated into protein only in male cells. By performing joint scRNA/ATACseq on *fruitless* neurons from both sexes (i.e. with and without Fru protein) we determine the transcriptional mechanisms by which Fruitless produces cell-type specific regulatory effects. We demonstrate that Fruitless’s actions are constrained by the chromatin landscape produced by cell lineage. Within lineage-specific landscapes, type-specific transcriptional dimorphisms arise from varying “Fru power” across cell types, which emerges as a combination of Fruitless dose, DNA binding domain isoforms, and the motif repertoire of individual enhancers. Together, these mechanisms allow a single factor to produce multifaceted transcriptional effects that subtly alter individual neuron types and produce sex-specific circuit functions.

## Introduction

Wiring a brain for behavior is a complex programmatic task: interconnected neurons have distinct cellular identities, are governed by distinct gene regulatory networks, and come from unrelated stem cell lineages (Tosches, 2017). Studies of innate behavior, and particularly innate sexual behavior, have given us strong insight into the gene regulatory networks which direct neuronal patterning. Neuronal development and circuit function at large are governed by hundreds of transcription factors (TFs) that collaborate in cell-type specific combinations, generating orders of magnitude more neuron types. Animals that reproduce sexually further diversify the circuits that can be built from a single genome via sex differentiation of select neuron types, which is implemented by master regulator transcription factors conditionally activated downstream of the sex-determination hierarchies (Gegenhuber et al., 2022; Roggenbuck et al., 2024; Sun & Tollkuhn, 2026). Sex differentiation thus provides a unique model to define how single transcription factors can work across diverse contexts to adjust precise aspects of neuronal development, including cell number, anatomy, connectivity, and physiology (Goodwin & Hobert, 2021).

Studies untangling the cell-type specific actions of TFs suggest specific regulatory outcomes can be conferred due to interactions with distinct sets of broadly expressed chromatin proteins and RNA processing factors (Carnesecchi et al., 2020); cooperative binding on DNA by interacting transcription factors, with motif spacing, orientation, and affinity heavily influencing gene regulatory outcomes (Luna-Zurita et al., 2016); and pre-existing differences in chromatin accessibility between cells allowing differential binding (Uyehara et al., 2022). These principles have not been empirically assessed in the context of neuronal diversity.

Among instinctual behaviors, the neural circuitry for courtship behavior in male fruit flies has been particularly well characterized. In this behavior, males integrate multisensory cues to detect a mate and then woo her by vibrating a wing to produce a courtship song. We know the cells that regulate and implement courtship, their developmental origins, their anatomic dimorphisms, and how they function at the circuit level. The patterning of courtship behavior is mediated by the male-specific <u>B</u>road-complex, <u>T</u>ramtrack and <u>B</u>ric-à-brac (BTB/POZ) family transcription factor Fruitless (Fru^M^) (Ito et al., 1996; Ryner et al., 1996). *fruitless* is transcribed from the *fru^P1^*promoter in 2-5% of neurons in the brains of both sexes (G. Lee et al., 2000).

There are at least 60 types of neurons within the *fru* set, including nearly all constituent neurons of the courtship circuit (Berg et al., 2025; Cachero et al., 2010; Deutsch et al., 2025; J. Y. Yu et al., 2010). Despite its transcription in analogous neuronal populations in both sexes, Fru^M^ proteins are only translated downstream of male sex determination programs, as the female-specific splicing factor Transformer (Tra) renders *fru*^P1^ transcripts untranslatable (Ito et al., 1996; Ryner et al., 1996). Translation of Fru^M^ is necessary in XY and sufficient in XX animals to generate courtship (Demir & Dickson, 2005; Ito et al., 1996). Fru^M^ shapes the final forms and circuit roles of post-mitotic neurons through the inhibition of cell death fates, alteration of axon and dendrite morphologies, and specification of synaptic connections. These effects are deployed in unique combinations across *fru* cell types (Berg et al., 2025; Cachero et al., 2010; Goto et al., 2011; Ito et al., 2016; Kimura et al., 2005; Kohl et al., 2013). While the circuit and cellular phenotypes generated by Fru^M^ have been extensively characterized, the developmental and molecular mechanisms by which Fru^M^ exerts these effects are poorly understood.

We and others recently described transcriptional mechanisms that connect neuronal lineage origins and mature circuit function in the *Drosophila melanogaster* cerebrum: Neurons derived from the same stem cell that share an anatomic blueprint (i.e. neurons of the same hemilineage) share a cocktail of about 10 hemilineage transcription factors; a separable set of birth order transcription factors diversify anatomy and connectivity within hemilineage patterning (Allen et al., 2026a; Cachero et al., 2025; Elkahlah et al., 2025). The cells that transcribe *fruitless* are specific birth order cohorts from within at least 60 of 180 hemilineage ground plans in the cerebrum (Cachero et al., 2010; Deutsch et al., 2025; H.-H. Yu et al., 2013). Thus, sex differentiated neurons do not share a single lineage origin, but instead are subsets of, and arise out of, many lineages (Allen et al., 2026b; Cachero et al., 2025; Elkahlah et al., 2025). The sex differentiation of each of these neuron types is thus just one component within each of their transcriptional recipes, and each time Fruitless acts, it does so in a distinct transcriptional context.

Fruitless is thought to act as a transcriptional repressor: Its best studied targets have lower expression when it is present; its motifs are found in enhancers that lose chromatin accessibility when it is present; and it interacts genetically and biochemically with heterochromatin-associated proteins (Brovkina et al., 2021; Ito et al., 2012, 2016; Meissner et al., 2016; Sato et al., 2020). While all the Fruitless protein produced in postmitotic neurons is the sex-differentiated Fru^M^ form (Brovkina et al., 2021; Sato & Yamamoto, 2022), sex-shared Fru^COMM^ is produced in embryogenesis and neurogenesis from promoters downstream of the sex-specific splicing site (Brovkina et al., 2021; Ryner et al., 1996; Sato & Yamamoto, 2022; Song et al., 2002); Fru^COMM^ has also been implicated in transcriptional repression (Calderon et al., 2022; Rajan et al., 2023). Fru^M^ proteins are expressed throughout metamorphosis (or pupation), when the larval brain and circuits are remodeled and built into the adult nervous system, and high Fru^M^ expression is maintained in the adult (G. Lee et al., 2000; Stockinger et al., 2005). Fru^M^ is necessary and sufficient for innate, female-directed courtship behavior in a critical window from ∼0-48h after puparium formation (APF); after this time, Fru^M^ remains important to restrict courtship to conspecific, virgin females (Arthur et al., 1998; J. Chen et al., 2021).4/14/26 11:04:00 AM

Here, we investigate the chromatin landscape upon which Fru^M^ acts, both over developmental time using bulk-ATAC-seq and at the cell-type level using joint scATACseq and scRNAseq. Using bulk ATAC-sequencing of Fru^M^ neurons and their female counterparts during developmental stages, we identify ∼2500 Fru^M^-closed regulatory regions. Instead of acting on a common set of effector genes to coordinate connections across layers of the courtship circuit, we determine that Fru^M^ works by subtraction: Overwhelmingly, we find that Fru^M^ results in enhancer closing at its DNA-binding motifs. By comparing male and female cells of matched lineage and birth order, we identify cell-type specific chromatin and transcriptional changes and interrogate their origins. We find that cell-type specific actions of Fru^M^ derive in part from lineage-specific enhancer opening, which constructs the chromatin landscapes on which Fru^M^ can act. In addition, different *fru* cell types exhibit strong and consistent variation in how impacted they are by Fru^M^. We find that this is influenced by overall *fru* dose and by the specific DNA binding domain repertoire of each cell type. Across cell types that vary in *fru* dose and isoform usage, closing of individual enhancers is influenced by their number, quality, and spacing of *fru* motifs. Together, this multitude of factors – lineage-patterned chromatin opening, transcription factor dose and isoform usage, and cis-element sequences – combine to allow cell-type specific gene regulation.

## Results

### When does Fru^M^ change chromatin accessibility during development?

In postmitotic neurons, transcription at the *fruitless* locus occurs from the P1 promoter, which produces the male-specific Fru^M^ isoform (Brovkina et al., 2021; Sato & Yamamoto, 2022). *fruP1* expression begins in late larvae, peaks in mid-pupal stages, and continues robustly in the adult; the effects of Fru^M^ — on chromatin, gene expression, circuit form, and circuit function – could arise at each of these stages. We previously identified differential chromatin accessibility in adult neurons downstream of Fru^M^ (Brovkina et al., 2021). To identify when in development Fru^M^ establishes sex-specific chromatin states, we extended our previous analysis in adults to pupal time points, 24 and 48 hours after puparium formation (APF). These timepoints encompass the highest levels of *fru* expression and overlap apoptotic events (12h-48h APF), neurite outgrowth (24-48h APF), initial formation of connections and synapses (48h+ APF), and the critical window for development of courtship behavior (Arthur et al., 1998; J. Chen et al., 2021; G. Lee et al., 2000).

Using fluorescence-activated cell sorting (FACS), we isolated a minimal population of central brain neurons expressing mCD8::GFP under the control of *fru^P1^*-GAL4 (“*fru* cells”) and performed the Assay for Transposase-Accessible Chromatin (Buenrostro et al., 2015) followed by high-throughput sequencing (ATAC-seq) on both *fru* and non-*fru* cells, as previously described (Brovkina et al., 2021) (≥3 biological replicates/stage, Figure 1A-C, Methods). Integrating these with our previous data from 2–7-day old adults, we identified regions whose accessibility correlated with the presence of Fru^M^ protein by comparing male *fruitless* neurons to the three other conditions (male non-*fru*, female *fru*, female non-*fru*) (Figure 1D,E). This approach successfully regresses out dosage compensation sites (Supplemental Figure 1F). We identified 3,138 (FDR < 0.01) Fru^M^-differentially accessible regions (Fru^M^-DARs) across time using DiffBind with a consensus peakset (Figure 1F, Supplemental Table 1). 578 regions were preferentially accessible in cells making Fru^M^ (Fru^M^-open regions), and 2,588 were closed in cells making Fru^M^ (Fru^M^-closed regions). While the ratio of Fru^M^-closed sites to Fru^M^-open sites is in concordance with our previous analysis, we recover more regions due to additional power and timepoints (Figure 1D, Supplemental Figure 1D, Methods). Among Fru^M^ DARs, some show differential accessibility across timepoints, while some do not (Figure 1G, Supplemental Figure 1G). None of these sites are detectable in simple male vs female comparisons, while 498 (∼16%) of these sites are differential in *fru* vs non-*fru* cell comparisons, suggesting population or cell-type specific regulatory regions (Supplemental Figure 1G).

**Figure 1.**
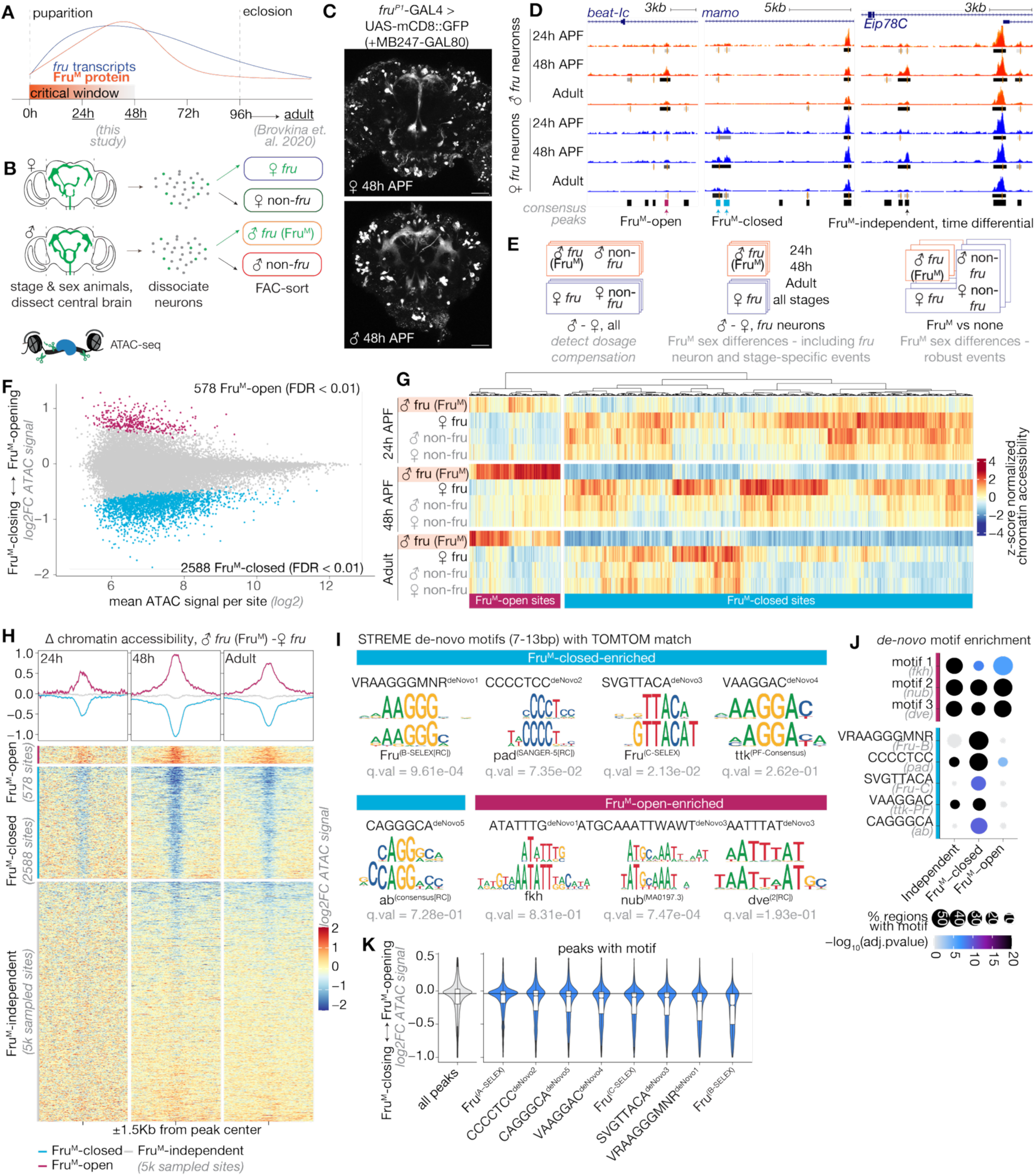
Fru^M^-dependent chromatin changes are instantiated by 48h APF and maintained in adults. A) Schematic of Fru^M^ levels and *fru* expression across pupal development, including the critical window for Fru^M^ function. Timepoints analyzed are underlined. B) Schematic of isolation method for *fru*-expressing neurons. C) 2-photon maximum intensity projections of GFP expression in sorted genotypes at 48h APF. D) Example UCSC genome browser screenshots of Fru^M^-dependent and independent chromatin changes across developmental time. Consensus peak bars below are determined by muMerge pooling of peaks shown under each condition. E) Schematic summarizing differential accessibility comparisons and their predicted detected events. F) MA-plot showing log2 baseMean chromatin accessibility of ∼41k consensus accessible peaks against log2 fold-change of neurons with Fru^M^ versus those without determined by DiffBind. Fru^M^-differentially accessible regions (Fru^M^-DARs) are colored. G) Heatmap displaying the relative changes in chromatin accessibility per Fru^M^-differential chromatin region across time and distinct sex, *fru*-expression, and developmental stage conditions. H) Signal heatmaps (DeepTools) of the difference in signal between pooled male and female *fru* conditions across Fru^M^-DAR regions and a 5K random sample of Fru^M^-independent regions. I) *De-novo* motifs found in Fru^M^-DARs. J) AME enrichment of *de-novo* motifs within open, closed, and independent sites; enrichment in each set calculated relative to shuffled control. K) Relative chromatin accessibility of peaks containing Fru motifs in Fru^M^ (i.e. male *fru*-expressing) neurons, arranged by mean.

As we observed previously, Fru^M^-DARs tend to be distal regulatory elements, with fewer than 5% of regions residing within 1kb of a transcriptional start site (TSS) (Supplemental Figure 1E). The genes nearest these DARs are enriched for synaptic, axon guidance, and cell adhesion functions, even when compared to all genes with accessible chromatin in developing, postmitotic neurons (Brovkina et al., 2021) (Supplemental Figure 1H). Some genes had multiple Fru^M^-DARs, suggesting highly cell-type specific regulation. Our consensus set of regions was called on all the timepoints together, and so we set out to determine their temporal dynamics. Fru^M^-directed opening and closing events called on the integrated dataset can be distinguished by eye in aggregate at 24h APF, but do not individually reach statistical significance (Supplemental Figure 1F), suggesting that the proportion of cells with remodeled events is not large enough statistically. Thus, chromatin remodeling is not yet temporally synchronized but has likely been initiated by 24h APF (Figure 1G, H). At 48h APF, we observe the greatest magnitude of differences in Fru^M^-DARs. Fru^M^-closed regions close gradually from 24h to 48h, and later from 48h to adult, such that the strongest closing signal per region is in the adult state (Supplemental Figure 2A, C). The ∼15% of regions that are detected only in male *fru* neurons (Fru^M^-open) emerge between 24 and 48h APF (Supplemental Figure 2A, B).

Female *fruitless* neurons, due to their transcription of *fru* but no translation of Fru^M^, are analogous to a *fru* knockout condition in our dataset; this allows us to analyze the dynamics of accessibility over developmental time for elements that Fru^M^ decommissions in male neurons. In female neurons (i.e. lacking Fru^M^), we find Fru^M^-closed regions to otherwise be accessible at 24h and increase in accessibility at 48h APF (Supplemental Figure 2). Most of the chromatin sites closed in male *fru* cells between 24h and 48h APF are maintained open in 2-7-day old adult female animals (Figure 1G, Supplemental Figure 2A, C). Others close in both sexes in adulthood, which is consistent with use of these genes in circuit development, which is mostly completed before adult eclosion (Supplemental Figure 2A, C) (Aimino et al., 2023; Muthukumar et al., 2014).

In addition to alternative promoter usage and sex-specific splicing at the 5’ end, *fruitless* transcripts also undergo mutually exclusive incorporation of terminal exons that encode distinct DNA binding domains. Three such protein isoforms, Fru-A, Fru-B, and Fru-C, are produced in postmitotic neurons and have different DNA binding motifs (Billeter et al., 2006; Dalton et al., 2013; Neville et al., 2014; von Philipsborn et al., 2014). To determine possible molecular drivers of differential accessibility, we performed formal motif enrichment analysis on Fru^M^-DARs as well as Fru^M^-independent sites using AME (MEME-suite, shuffled background). Amongst the top 15 enriched motifs for each set, we found motifs for Trl (GAGA factor) and Klu (Supplemental Figure 2D), whose motifs have been suggested to be a general feature of neuronal enhancers (Coyne et al., 2025). As we did in adults, we again find the canonical Fru-B motif (Fru-B^SELEX^) to be highly specifically enriched in Fru^M^-closed regions (Figure 1I, J, Supplemental Figure 2D)(Brovkina et al., 2021). Because this method is limited to known PWMs, we further performed *de novo* motif analysis using STREME (7-13bp). Our top *de novo* motif is essentially identical to the known SELEX motif for Fru-B (Figure 1I, Supplemental Figure 2F). We also identify three motifs with similarity to Fru-B but matching closer to known motifs for Pad (CCCCTCC^deNovo2^), Ttk-PF (VAAGGAV^deNovo4^), and Ab (CAGGGCA ^deNovo5^). Ttk and Ab are BTB-TFs whose zinc fingers are closely related to each other and to those of Fru (Bonchuk et al., 2024; Neville et al., 2014).

Lastly, we identify a de-novo motif representing a more minimal sequence within the canonical Fru-C motif (Fru-C^SELEX^). Given that we do not find Fru-C^SELEX^ enriched in Fru^M^-closed regions, this *de novo* motif may more accurately reflect Fru-C binding *in vivo*. We did not identify motifs resembling canonical Fru-A (Fru-A^SELEX^) motifs in either analysis. Interestingly, while Fru-C^SELEX^ and Fru-A^SELEX^ resemble one another, the *de novo* motif we find for Fru-C is more distant from Fru-A^SELEX^ (Supplemental Figure 2F). We formally measured enrichment of these *de-novo* motifs using AME and find very specific enrichment of all motifs except the Pad-like CCCCTCC motif in Fru^M^-closed regions (Fig 1J). Loosening the threshold for calling a peak “Fru^M^-closed” to FDR < 0.05 had nearly no effect on the set of *de novo* motifs discovered nor their order of enrichment, suggesting that detection of these motifs is robust (Supplemental Figure 2E). Motifs in the Fru-B and Fru-C families have the strongest impact on chromatin closing Fru^M^ cells (Fig 1K).

### Cell-type specific profiling of chromatin accessibility and gene expression

Studies of constituent neurons in the *fruitless* circuit found major morphological groups that derived from at least 60 distinct stem cell lineage origins (Cachero et al., 2010; J. Y. Yu et al., 2010). Individual *fru* neuron types perform different circuit roles and are differentiated in distinct ways (Cachero et al., 2010; Ren et al., 2016; H.-H. Yu et al., 2013; J. Y. Yu et al., 2010). Our previous analysis suggested that Fru^M^-DARs are cell-type specific (Brovkina et al., 2021). To test this model at scale, identify regulatory mechanisms, and link chromatin accessibility to gene expression, we jointly measured chromatin accessibility (scATAC-seq) and gene expression (scRNA-seq) using 10X Multiomics on single cells. Due to the cumulative nature of chromatin accessibility, the endurance of earlier developmental events into adult neurons (Fig 1G,H, Supplemental Figure 3A-C), and the technical complexity of multiomics in *Drosophila* neurons, we profiled the adult stage.

We FAC-sorted *fru* neurons using a similar genotype and methodology as our bulk sequencing and collected five independently sorted libraries (Figure 2A, Supplemental Figure 1C). Three were mixed-sex to minimize confounding batch effects with sex differences, and two were split-sex to validate our post-hoc determination of cell sex. To maximize coverage of rare cerebral cell types, we reduced the stoichiometry of the optic lobes in our dissections. After QC (Supplemental Figure 3A-G), we obtained multiomics profiles for 20,719 cells, representing ∼10-fold coverage of *fru* neurons in the central brain (i.e. 5-fold in each sex). All cells expressed high levels of *fru* and the neuronal marker *nSyb*; GAL4 and mCD8::GFP transcripts, while sparser, follow *fru* expression (Supplemental Figure 3H).

**Figure 2.**
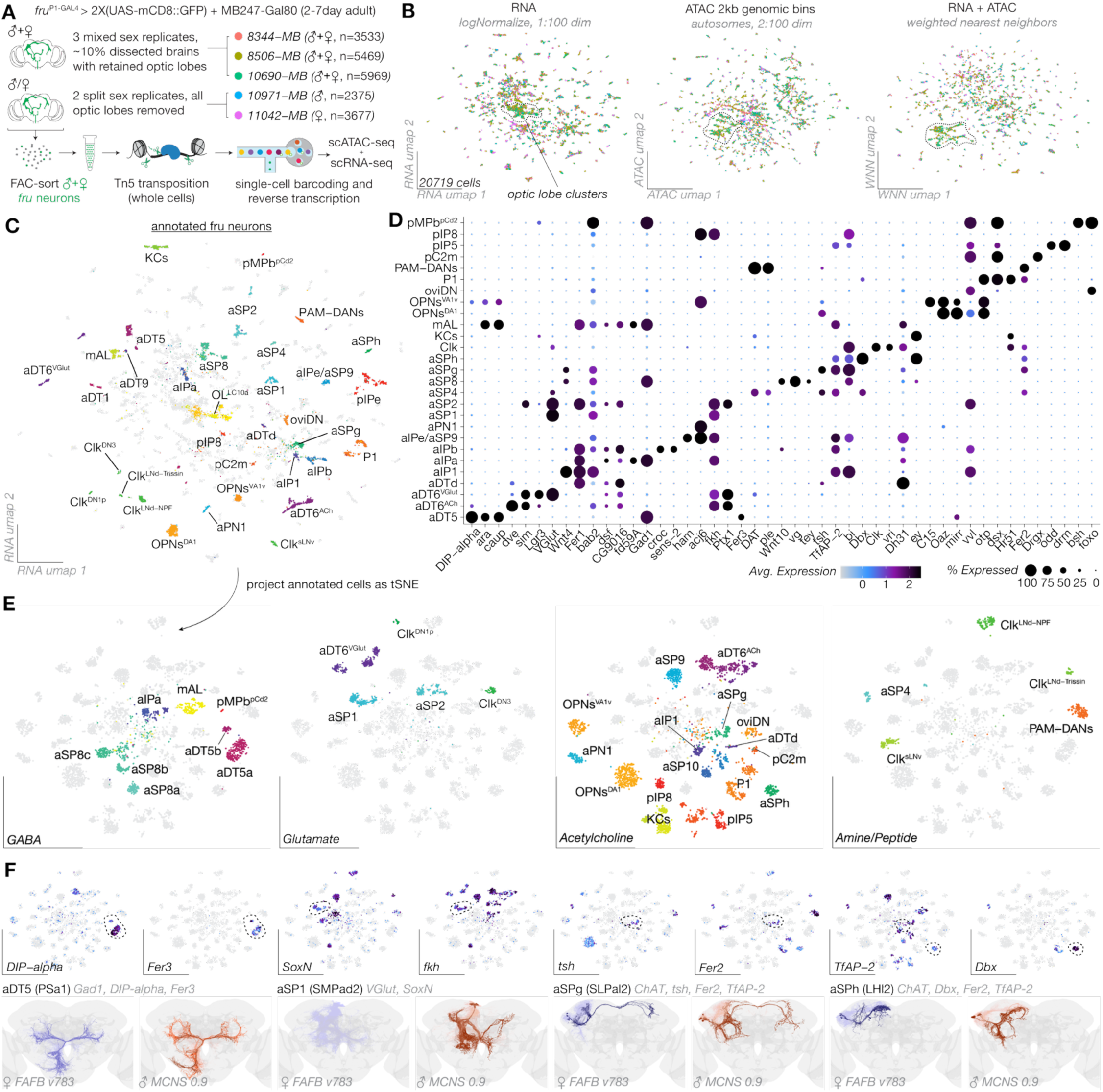
Single-cell multiomic analysis of adult *fru*-neurons. A) Schematic of experimental paradigm for isolating *fru* neurons showing independent library collections and number of final cells passing QC per replicate. B) UMAP dimensionality reductions made on RNA, ATAC (2kb genomic bins), and weighted nearest neighbor analysis between RNA and ATAC modalities C) RNA modality UMAP projection of *fru* neurons, with annotated neurons highlighted. D) Dotplot of known and discovered marker genes for each group of annotated neurons in Fig 2C. E) tSNE RNA projection of annotated neurons subset and reclustered from full multiome data, split by neurotransmitter class. F) Markers of newly annotated *fru* neuron types including aDT5, aSP1, aSPg, and aSPh (top) with reconstructed EM skeletons of neurons from both female (FAFB v783) and male (MCNS 0.9) neurons shown; whole hemilineages are shown in lightest colors, with annotated *fru* neurons in darker colors.

We used ATAC signal across genomic tiles and normalized gene expression to perform weighted-nearest neighbor cell clustering without prior expectation (Figure 2B), producing ∼220 groups of cells distinguishing anatomically well-studied and lineage-defined *fru* cell types (Figure 2C). More numerous canonical types split into multiple clusters along expression of TFs we recently reported as reflecting birth order within lineages (Elkahlah et al., 2025). Indeed, we and others have previously defined birth-order-related transcriptional subtypes of aDT6, P1, and mAL neurons, e.g. mAL^Br^, mAL^pdm3^ (Allen et al., 2026b; Elkahlah et al., 2025). Morphological and connectivity subtypes within lineage-defined *fru* classes are also visible in anatomic studies, with the degree of diversity varying according to how “cell type” is defined (Costa et al., 2016; G. M. Rubin et al., 2025). At this resolution, cell types did not split by sex, consistent with recent analyses that place sex as a subsidiary axis within lineage and birth order patterning (Allen et al., 2026a; Elkahlah et al., 2025; Cachero et al., 2025).

Using all known previously annotated markers of *fru* neuron subtypes (Supplemental Table 3), we assigned 51 single-cell clusters to 31 anatomic types of *fru* neurons from the cerebrum and optic lobe (Figure 2C-E). Discriminating markers across annotated types include primarily hemilineage transcription factors (Elkahlah et al., 2025), functional/state TFs (*Clk*, *vri*, *dsx*), neurotransmitter-associated genes, morphogens, and peptides (Figure 2D,E, Supplemental Table 4), consistent with the different lineage origins of *fruitless* neurons and their functional diversity. Of these annotated types, many groups had not been transcriptionally defined previously. For example, we were able to identify aDT5, which is part of the PSa1_lateral hemilineage, due to *DIP-alpha* expression (Palmateer et al., 2023), GABAergic identity (Eckstein et al., 2024), and the hemilineage transcription factor Fer3 (Elkahlah et al., 2025) (Figure 2E,F). aSP1 neurons (SMPad2 hemilineage) are defined by expression of *NP111*^SoxN^-Gal4 (J. Y. Yu et al., 2010) and VGlut-Gal4 (von Philipsborn et al., 2014) (Figure 2E,F). We identified aSPg (SLPal2 hemilineage) and aSPh (LHl2_dorsal hemilineage), both of which are predicted cholinergic in the connectome, using a Fer2-TfAP2 split-GAL4 (Yunzhi Lin, manuscript in preparation). We discriminated between the two using *tsh*, which was previously shown to label aSPg (also known as aSP6) but not aSPh neurons (J. Y. Yu et al., 2010). We used similar methodology to annotate other *fru* neuron types (Supplemental Table 3). For concision, we focus most subsequent analyses on this annotated subset (Figure 2E).

### Chromatin landscapes underlying distinct fru neuronal cell types

We previously validated cell-type specific enhancers identified within the *fru* locus from bulk ATAC-seq, showing that they drive reporter gene expression in different sets of *fru* neurons (Brovkina et al., 2021). Generating single-cell ATAC coverage tracks for the *fru* gene locus, we indeed find distinct patterns of chromatin accessibility across *fru* neuron types (Figure 3A), and matched enhancer-reporters drive expression in the expected cell types (Jenett et al., 2012; Pfeiffer et al., 2008) (Figure 3B). Thus, our scATAC data and cluster annotations can give us real insight into cell-type specific enhancer usage, and further, shows distinct chromatin accessibility profiles between *fru* neuron types.

**Figure 3.**
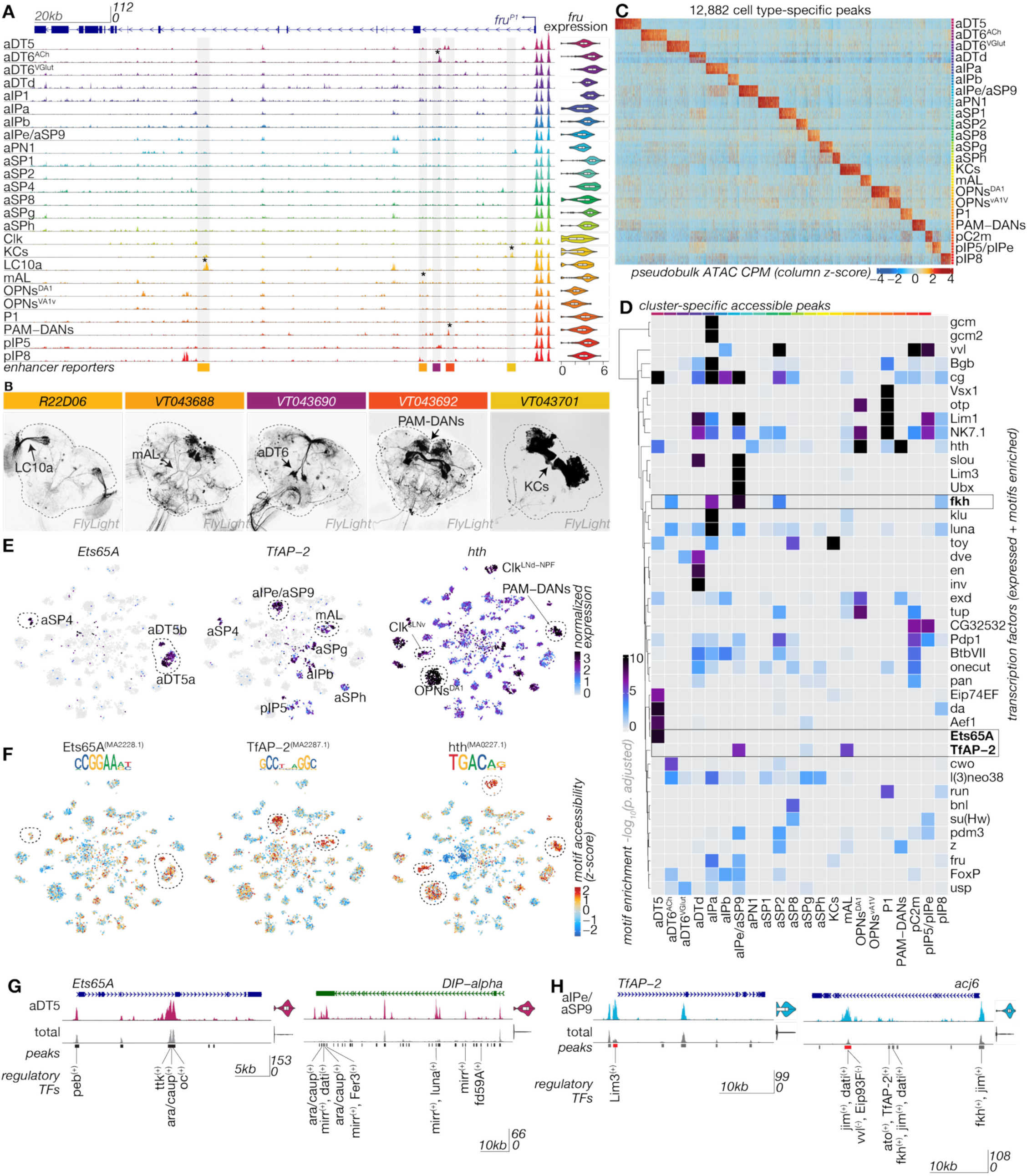
Chromatin landscapes underlying distinct *fru* neuronal cell types. A) Coverage plot of cell-type pseudobulk ATAC signal across the *fru* gene locus per selected annotated *fru* neuron types. Highlighted regions correspond to enhancer tiles in (B) in left-right order. C) Heatmap of marker peaks per annotated cell type; replicate and cell-type pseudobulk ATAC signal. Male and female separated libraries are merged into one replicate to match the others. D) Heatmap of adjusted p-values for enrichment of transcription factor motifs in significant marker peaks for each annotated cluster. TFs not expressed in a cell type are zeroed. E) Expression of three transcription factors across annotated cells and (F) chromVAR deviation scores for Ets65A, TfAP-2, and Hth motifs representing a bias-corrected measurement of the accessibility of regions with a motif across individual cell types. G-H) ATAC coverage tracks of example cell-type marker genes with cell-type specific enhancers annotated with PANDO predicted regulatory TFs in (G) aDT5 neurons and (H) aIPe/aSP9 neurons.

Annotated *fru* neuron types derive from different hemilineages and represent specific birth order cohorts; most of the chromatin landscape of each cell type will reflect these axes, with sex-specific events a minor contributor (Allen et al., 2026a,b; Cachero et al., 2010; Elkahlah et al., 2025). We first explored these aspects of neuronal chromatin, independent of sex or Fru^M^. We used our consensus peakset to find enriched peaks for all clusters, then took the top thousand marker peaks in annotated neuron types (Figure 3C, Supplemental Figure 1A). To determine the transcription factors positively regulating cell-type specific enhancers, we identified marker peaks (p.adj < 0.001) for annotated neuron types and performed motif enrichment analysis (Figure 3D, Supplemental Table 7). Because many transcription factors share motifs, particularly homeodomain TFs that mark hemilineages, we only retained motifs for TFs with expression in that cell type; motif enrichments unfiltered by TF expression are in Figure S3C. This represents a set of likely activators of gene expression and chromatin accessibility within these cell types (Supplemental Table 7). Most of these TFs have been previously annotated as hemilineage or, more rarely, birth-order TFs (Allen et al., 2026a; Cachero et al., 2025; Elkahlah et al., 2025).

Multiomic data gives us the unique ability to relate transcription factor expression, motif presence, and chromatin accessibility to identify potential molecular mechanisms underlying the activity of a given transcription factor (Figure 3E, F). For example, *Ets65A*, *TfAP-2* and *Hth* RNA each positively predicted opening of their corresponding motifs (Supplemental Figure 3D, Supplemental Table 6). Ets65A presence explained a subset of cases in which its motif opened, with motif opening in other cell types likely due to other Ets family members. TfAP-2 motifs opened in only a subset of cell types with *TfAP-2* expression, suggesting possible combinatorial relationships with other factors.

We used chromVAR to calculate a per-cell chromatin accessibility score (bias-corrected “deviation”) of peaks associated with specific motifs, relative to the expected chromatin accessibility across all cells (Schep et al., 2017); positive scores represent relatively accessible motifs, while negative scores represent relatively inaccessible motifs in a given cell. By correlating the expression of a TF with the chromVAR deviation score of its motifs on a per-cell basis (Calderon et al., 2022), we predict activating transcription factors across the dataset, as well as putative repressors (Supplemental Figure 4D, E). Amongst the top TFs correlated with open chromatin in the dataset, we find *rn, CG443869,* and *kn* (Supplemental Figure 4F). TFs whose expression is most strongly correlated with chromatin closing include *lov, Eip93F*, and *pdm2* (Supplemental Figure 4G). Notably, many of the top activating TFs are associated with hemilineage identity, while the top repressive TFs are associated with birth order.

Many TFs reflecting hemilineage and birth order have enduring expression from early neuronal specification and are maintained into the adult (Elkahlah et al., 2025). To determine how the transcriptional networks underlying neuronal fate are stably maintained and induce expression of target genes, we ran PANDO for gene regulatory network inference (Fleck et al., 2023). We identified many cases in which hemilineage transcription factors are predicted to regulate each other, as well as non-TF genes important for neuronal features. For example, Ets65A is a characteristic hemilineage factor expressed in aDT5 neurons, of the PSa1_lateral hemilineage. An enhancer in *Ets65A* that is specifically accessible in those cells is putatively regulated by other hemilineage transcription factors, Ara and Caup (Figure 3G; Ara and Caup share a motif and are usually co-expressed). *DIP-alpha*, a neuronal adhesion gene that is a marker of aDT5 neurons, contains enhancers predicted to be regulated by hemilineage factors Ara, Caup, Mirr, and Fer3 (Figure 3G). In aIPe/aSP9 neurons, which are from a different hemilineage (VLPl2_posterior), we again find predicted cross-regulation among hemilineage TFs, with Lim3 predicted to induce *TfAP-2* (Figure 3H), TfAP-2 and Fkh predicted to induce *acj6*, and Vvl predicted to repress *acj6* (Figure 3H). Notably, *vvl* and *acj6* have mutually exclusive expression patterns elsewhere in the brain (Komiyama et al., 2003; Li et al., 2017). In addition to hemilineage TFs, we also see birth-order TFs such as Jim, Dati, and Eip93F putatively regulating *TfAP-2* and *acj6*. Lastly, looking at the timing of accessibility of these cell-type specific enhancers, we find that they are not universally opened in neuronal precursors (Lucas et al., 2021), but we see established accessibility in post-mitotic *fru* neurons by 24h APF (Supplemental Figure 4H).

### Adult sex differences in gene expression in fru neurons

Having identified that Fru^M^ is a repressor and the lineage and birth order-specific chromatin accessibility landscape that it can act on, we next went on to ask about cell-type specific sex differentiation. We used *rox1* and *rox2* RNA and X::autosome ratio in DNA reads to call cell sex in mixed-sex libraries and benchmarked these assignments using single-sex libraries (Figure 4A,B, Methods). Some clusters, such as *dsx*-expressing P1, pC2l, and pC2d, contained only male cells, as expected (C. Zhou et al., 2014). Others comprise relatively even numbers of male and female cells, and a few are enriched for female cells. For each cell type, we calculated the ratio of male to female cells and delineate between cell types with sex-shared counterparts and those which appear to be male-specific and treat these cell types distinctly (Figure 4 C,D, Supplemental Figure 5A-C). Female-specific cell types are individually rare and are under-annotated both in our cell set and in the field (Allen et al., 2026b).

**Figure 4.**
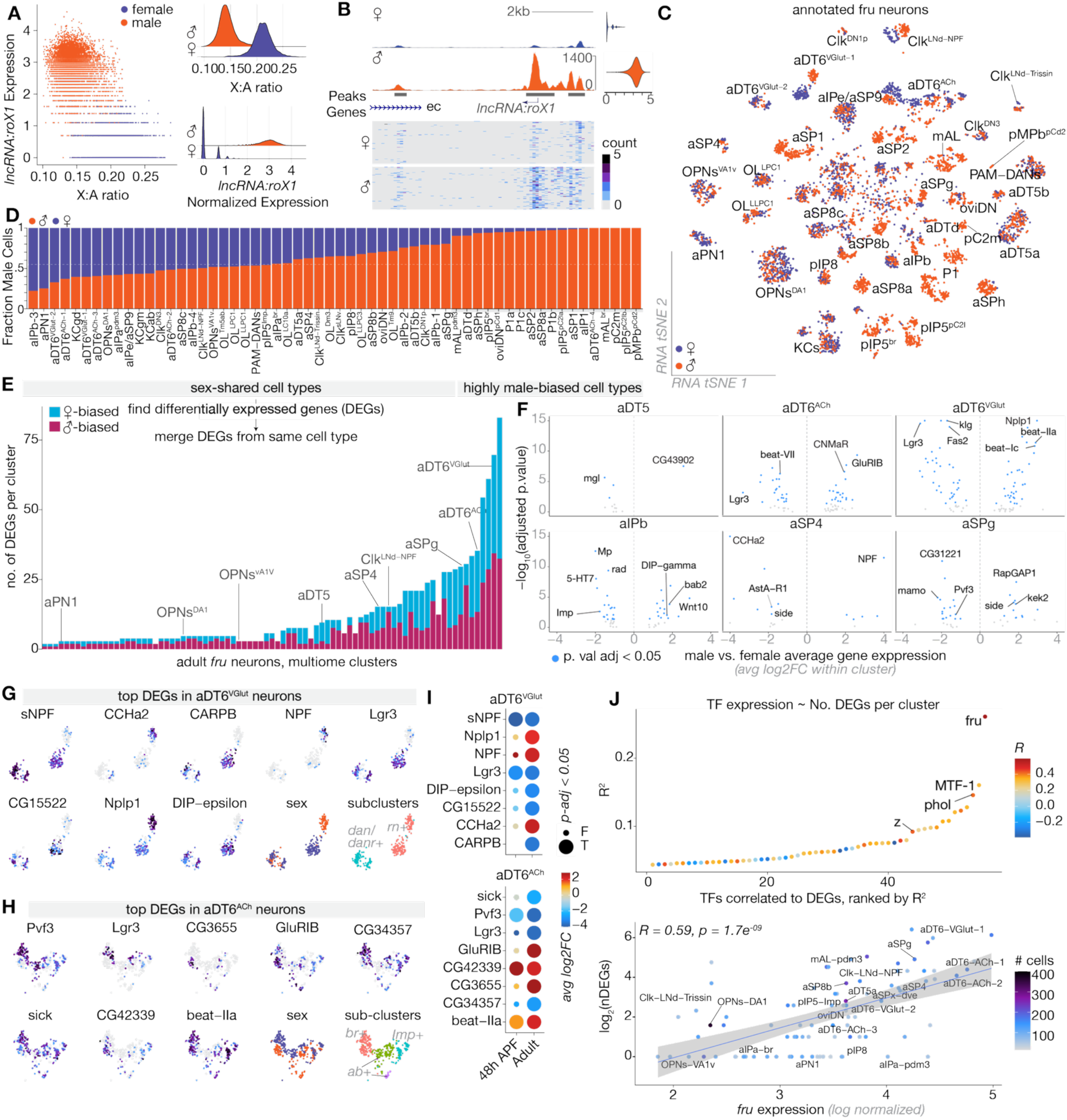
Adult sex differences in gene expression in *fru* neurons. A) X:Autosome ratios of chromatin accessibility counts against male-specific *lncRNA:roX1* expression in demixed male and female *fru* neurons. B) Coverage plot of *lncRNA:roX1* genomic locus in demixed neurons with cell sample of counts. C) RNA tSNE projection of subset annotated clusters, colored by sex. D) Sex ratios of all annotated cell types in adult multiome data, ordered by percent male cells and with total percent in dataset shown by dashed line. E) Number of differentially expressed genes between male and female neurons in sex-shared cell types by DESeq2 on pseudobulked replicates. F) Volcano plot of sex differences in gene expression in select *fru* cell types; most significantly changed genes labeled. G-H) RNA tSNE of top 8 differentially expressed genes, assigned sex, and subcluster identity based on top differentiating genes in (G) aDT6-VGlut neurons and (H) aDT6-ACh neurons I) Male vs female average log2 fold change and significance of genes in G and H in adult multiome data and matched cells from Palmateer 2023. J) Top: Number of DEGs per cluster correlated to expression of 280 expressed TFs, ranked by R^2^. Bottom: Correlation of DEGs (log2) per cluster to log normalized *fru* expression. R value and p-value of correlation are reported. Points are colored by number of cells per tested cluster.

To determine transcriptional sex differences, we took advantage of our four independent libraries per sex, pseudobulked cells per cluster, and performed DESeq2 differential gene expression, which results in robust differential expression testing (Supplemental Figure 5D). Because sex differentiation occurs within specific birth order cohorts within a hemilineage (Elkahlah et al., 2025), we performed differential gene expression testing on a more granular cluster level where cell types were split by birth order into distinct clusters; differentially expressed genes from within birth order cohorts were then aggregated at the hemilineage level. This avoids confounding differential expression related to birth order TFs with differential expression related to sex.

For cell types present in both sexes, there is a broad range in the degree of transcriptional sex differences (Figure 4E,F, Supplemental Figure 5G). After removing genes that are themselves part of the sex differentiation and dosage compensation cascade (*rox1, rox2, Sxl,* and *ssx*), many clusters show zero transcriptional sex differences. These include previously well studied *fru*-expressing cell types such as Kenyon cells, which have no known anatomic or functional sex differences; and olfactory projection neurons, which have differential connectivity between the sexes that does not depend on cell-autonomous Fru^M^ (Figure 4E, F) (Kohl et al., 2013). The majority of sex DEGs are only differentially expressed in one cell type, though others are observed multiple times (Supplemental Figure 5E, Supplemental Table 8). Many of them fall into the category of “cell surface and secreted molecules,” especially IgSF domain proteins.

To better understand the transcriptional effects of Fru^M^, we focused on two cell types – aDT6^VGlut^ and aDT6^Ach^ neurons, which arise from the FLAa3 hemilineage and are thought to be important for maintaining the precise sequence of courtship steps (Figure 4G) (Allen et al., 2026b; Manoli & Baker, 2004). These cells are present in both sexes and anatomically similar (Berg et al., 2025), yet have among the highest number of sex DEGs. We observe five birth order cohorts of *fru* aDT6 neurons, with distinct outputs of Fru^M^: The glutamatergic, *rn*-expressing and cholinergic, *Br*-expressing birth order cohorts have strong transcriptional sex differences that are distinct from one another. These groups separate by sex in UMAP and tSNE space. The other birth order cohorts have minimal transcriptional sex differences. The top differential genes in adult aDT6 neurons show the same direction and often comparable magnitudes of gene expression changes in development (Figure 4H) (Palmateer et al., 2023). While we cannot identify sex-differential gene expression differences in bulk RNA-seq data from *fru* neurons due to cell-type specificity, we find sex-differential genes across clusters to be used dynamically across neuronal maturation in the pupal stage (Supplemental Figure 5F).

Lastly, we sought explanatory variables for the magnitude of sex differences in gene expression across cell types. The number of sex-DEGs was not correlated with the number of cells per type (Supplemental Figure 5G). For each sex-shared cluster in the data, we asked whether any transcription factor’s expression pattern predicted the number of sex-DEGs. We find that by far the strongest predictor of the number of sex-DEGs per cell type is the expression level of *fru* itself (Figure 4I). Strikingly, *fru* expression varies ∼20-fold across its expressing cell types (i.e. expression between e^2^ and e^5^ in Figure 4J, Supplemental Figure 3H).

### Cell-type specific changes in chromatin accessibility

We next sought to understand the Fru^M^-dependent chromatin changes that underlie cell-type specific sex differentiation events. Fru^M^ is known to have two developmental roles: It both shapes the stoichiometry of male versus female *fru* cell types, mostly by suppressing lineage-programmed apoptosis in male cells (Kimura et al., 2005), and adjusts transcription in male *fru* cells. Because Fru motifs are present only in regions closed in the presence of Fru^M^, we previously hypothesized that these are Fru’s direct targets, while “Fru^M^-open” regions are lineage-specific enhancers related to cell identity independent of Fru^M^ – i.e. cells absent from our population of female *fru* neurons (Brovkina et al., 2021) (Figure 1). To test this hypothesis, we took regions we annotated as Fru^M^-open and Fru^M^-closed in our bulk ATAC dataset and asked which cell types contributed to the accessibility of these regions (Figure 5A).

**Figure 5.**
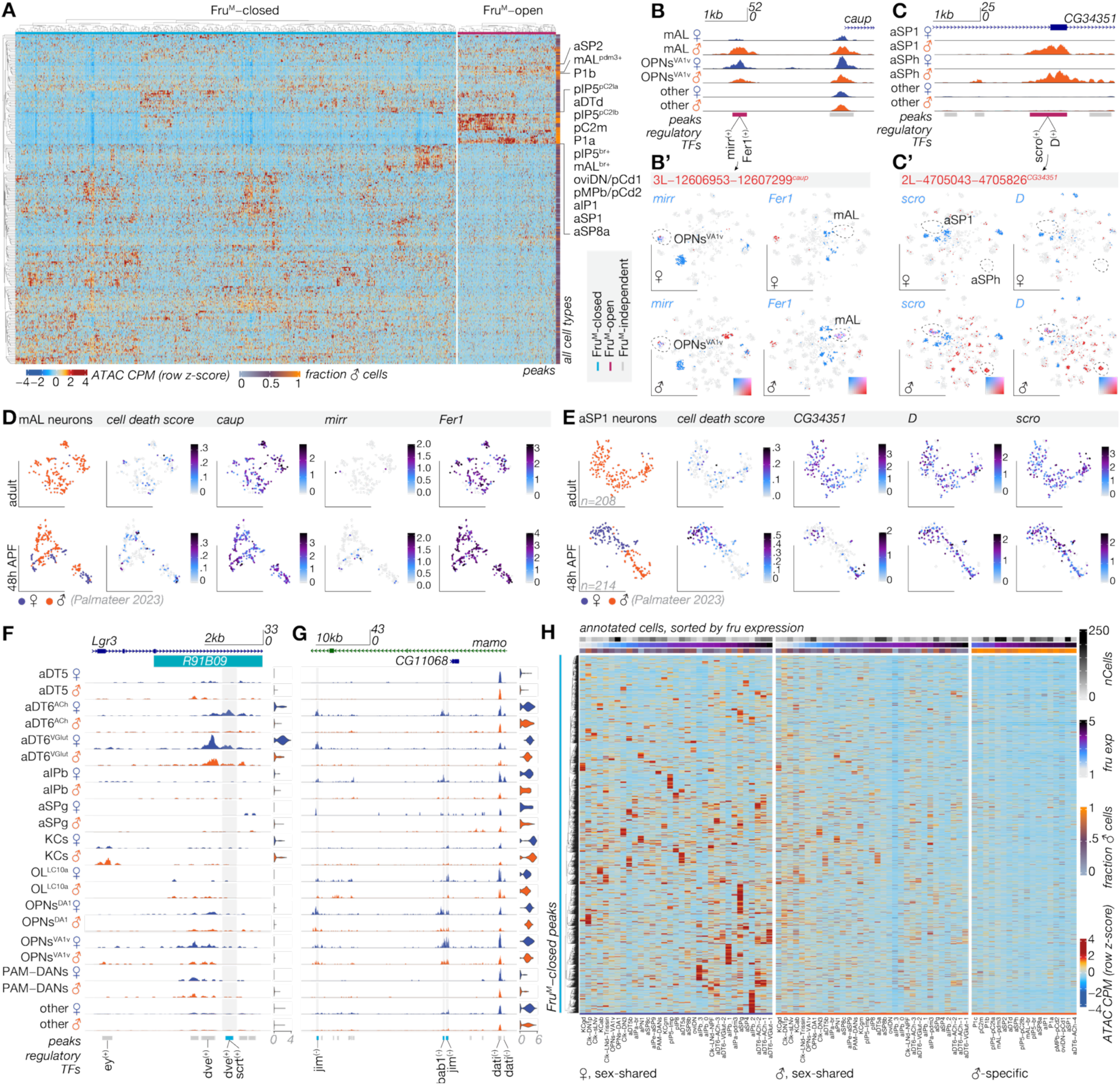
Cellular events underlying changes in chromatin accessibility in Fru^M^ neurons. A) Heatmap of ATAC CPM across identified Fru^M^-DARs from bulk ATAC-seq data. Notable male-specific or highly male-biased cell types are labeled. B) Fru^M^-open region upstream of *caup* mapped to mAL neurons and VA1v OPNs with PANDO predicted regulatory TFs shown. B’) Expression of PANDO predicted TFs *mirr* and *Fer1* (blue) overlapped with enhancer accessibility (red) in annotated cell types. C) Fru^M^-open region in *CG34351* mapped to aSP1and aSPh neurons with PANDO predicted regulatory TFs shown. B’) Expression of PANDO predicted TFs *D* and *scro* (blue) overlapped with enhancer accessibility (red) in annotated cell types. D-E) UCell score of cell death genes (*grim*, *rpr*, *hid*, *skl*), and expression of genes from (B) in mAL neurons (D) and (C) in aSP1 neurons (E) in adult multiome data (top) and 48h APF scRNAseq data from Palmateer 2023 (bottom). F-G) ATAC coverage plots of annotated *fru* cell types split by sex with PANDO predicted regulatory linkages shown. In (F), cell-type specifically accessible Fru^M^-closed region in *Lgr3* highlighted; in (G), Fru^M^-closed enhancers in *mamo* opened in multiple female cell types. H) Heatmap of chromatin accessibility of all Fru^M^-closed regions across annotated cell types, ordered by *fru* expression. For sex-shared neurons, same ordering is used for male and female cells.

The differential accessibility of Fru^M^-closed regions emerged throughout the cell-type repertoire. In contrast, we find that the accessibility of Fru^M^-open regions is strongly driven by cell types with high male::female ratio. For example, we identified a Fru^M^-open element upstream of *caup*, a hemilineage TF of CREa1B (mAL neurons) and ALad1B (VA1v olfactory PNs) (Figure 5B). we find this element to be accessible in male mAL neurons, which are far more numerous than their female counterparts, but equally open in olfactory PNs, which have a balanced sex ratio. This enhancer is predicted to be positively regulated by hemilineage factor Fer1 in CREa1B and Mirr in ALad1B and has no *fru* motifs. Similarly, we find a Fru^M^-open element in *CG34351*, accessible in male-specific aSP1 and aSPh neurons. This element is predicted to be positively regulated by Scro and D and has no *fru* motifs (Figure 5C). Scro and DD are both expressed in aSP1 neurons; other factors must open or maintain this region in aSPh neurons.

If these enhancers and their downstream genes are indirectly sex-differential due to cell death, a timepoint when mAL neurons and aSP1 neurons are present in the female should show equal expression of the enhancer-regulated genes and their upstream regulatory transcription factors. AtAt 48h APF (Palmateer et al., 2023), we can see a greater proportion of female mAL neurons and balanced male and female aSP1 neurons (Figure 5D, E). Indeed, in female mALs, *caup* and *Fer1* are expressed along with apoptosis-promoting transcripts, indicating that the female neurons have not completely undergone cell death. Strikingly, aSP1 neurons show expression of cell death genes only in female neurons, while the expression of *CG34351, scro,* and *D* are equal (Figure 5E). These data demonstrate that “Fru^M^-open” regions are lineage-specific enhancers in cells with a higher stoichiometry in male. Their sex bias in bulk data results from the impact of Fru^M^ on cell type distribution. While we were able to anticipate this result due to the field’s dense knowledge of sex-differentiated cell type distribution in this system, we belabor the point for two reasons: First, positive assignment of the source of these DARs allows us to rule out alternate models (e.g. that Fru^M^ moonlights as an activator; can interact with enhancers lacking Fru^M^ motifs through a cofactor; or has a substantial activating role through disinhibition). Second, these results allow us to emphasize that differences in cell fates may confound *any* studies of state-dependent gene regulation that lack anatomic ground truth, whether these rely on bulk sequencing or on incompletely resolved scRNAseq clusters.

We next focused on determining the cellular source of Fru^M^-closed events. As we showed in Figure 1, these regions contain Fru^M^ motifs at a high rate. We sought to ask two questions about the origin and regulation of Fru^M^-regulated enhancers: 1) were these sites opened in cell-type specific or universal ways, and 2) when these regions are closed, are they closed across all cells in which they had been opened? Despite the immense complexity of the transcriptional landscape, we can link male-repressed genes to specific Fru^M^ chromatin-closing events and assign them to known neuron types. As an example of a Fru^M^-closed region that is open cell-type specifically, we use the validated enhancer in *Lgr3*, which we find accessible only in female aDT6 neurons (Figure 5F). This enhancer is predicted to be positively regulated by Dve, which is a marker TF for this hemilineage (Allen et al., 2026a,b). *Lgr3* is also robustly repressed in expression in male aDT6 neurons, as has been shown using gene reporters previously (Meissner et al., 2016). We reason that this enhancer was opened selectively in this lineage during development and later closed in male cells by Fru^M^. This example represents the more common statistical class: Fru^M^-closed regions tend to be cell-type specifically accessible in females rather than universally accessible across all *fru*-expressing cells, suggesting that their chromatin accessibility is already lineage-specific prior to Fru^M^ activity (Figure 5H).

Still, cases where Fru^M^-closed regions are accessible across many cell types in females do exist, such as a regulatory region of the transcription factor *mamo* that is open in many female neurons types and closed in most of their male counterparts (Figure 5G). We reason that enhancers following this pattern were opened in many lineages and later closed by Fru^M^ across multiple male cell types. Accessibility within birth-order defining transcription factor loci contributes a large amount of variation to the ATAC landscape of our dataset overall, defining principal component 2 (Supplemental Figure 6B). While much of this variation is produced by birth order diversification itself (i.e., upstream of *fru*), these factors are also a common class of predicted Fru^M^ targets. Fru has previously been shown to bind such loci (Neville et al., 2014). We find regulatory elements fated to be closed by Fru^M^ in postmitotic neurons to acquire accessibility earlier in neuronal development, with subsets becoming accessible in Type I neuroblasts in the embryo, Type II neuroblasts in L3 larvae, and postmitotic neurons in L3 larvae; these likely represent more commonly commissioned regulatory regions across neurons (Supplemental Figure 6C).

Towards the cell-type specific effects of Fru^M^, we also find that regions with the potential to be closed by Fru^M^ (i.e. that close in some male cell types) can remain accessible in other cell types – i.e. the strength of Fru^M^-closing is also heterogeneous across cell types. For example, KCγ’s and aDT5 neurons show weak closing, while others such as the aDT6 populations have strong differences, with few Fru^M^-closable sites remaining accessible in male neurons (Figure 5H, Supplemental Figure 6D). While cell number could bias the detection of Fru^M^-closed regions in bulk data, the latter cell types are comparable in number. Much like how *fru* expression partially predicts the number of sex-DEGS, cell types with higher *fru* dosage show very little accessibility across Fru^M^-closable sites (Figure 5H, Supplemental Figure 6D). Further, we find that male-specific neuron types have among the highest *fru* expression and the least accessibility across bulk Fru^M^-closed regions (Figure 5H). While calling DARs based on the bulk dataset risks under-sampling enhancers that are closed in rare cells, our gene expression analysis (Figure 4E, Supplemental Figure 5G) suggests that this is not the primary variable influencing the degree of observed sex difference.

In male-specific cell types, it is possible that Fru^M^-closed regions were simply never accessible, as we lack female cells with which to compare them. However, the preponderance of Fru^M^-motifs in these regions suggests the hypothesis that cells subject to the strongest sex differences via apoptotic escape are also subject to strong Fru^M^ closing at other enhancers – i.e. it appears to take a lot of Fru^M^ to survive lineage-programmed apoptosis and this dose robustly closes any open enhancers with Fru^M^ motifs (Supplemental Figure 6E). We therefore find two explanatory variables for cell-type specific sex differences: the landscape that Fru^M^ has access to is cell-type specific, and its potency to modify these landscapes is highly variable.

### Dose and isoform usage of Fru^M^ across unique sequence landscapes

To test directly whether increasing Fru^M^ dosage correlates with stronger chromatin closing effects, we assessed how *fru* expression related to Fru^M^ motif accessibility across individual neurons (Figure 6A, B). We find significant anticorrelation between *fru* expression and the accessibility of all SELEX and *de novo* motifs. The strongest relationship between dose and closing were for Fru-B-like motifs (especially VRAAGGGMNR^deNovo1^), followed by Fru-C motifs, and a slight effect of Fru-A (Figure 6A, B, Supplemental Figure 7A, B). In female cells, we fail to find a relationship between *fru* expression levels and motif accessibility across any motif, since female cells lack Fru^M^ protein. Across all transcription factors with known PWMs, the only TFs with differing modes of action between male and female neurons are Fru^M^ motifs and Dsx motifs, suggesting that these sex-differentiation factors are not globally modifying the activity of other transcription factors across the data (Figure 6C), though Fru^M^ could still produce sex differentiated activity of downstream TFs in specific cell types.

**Figure 6.**
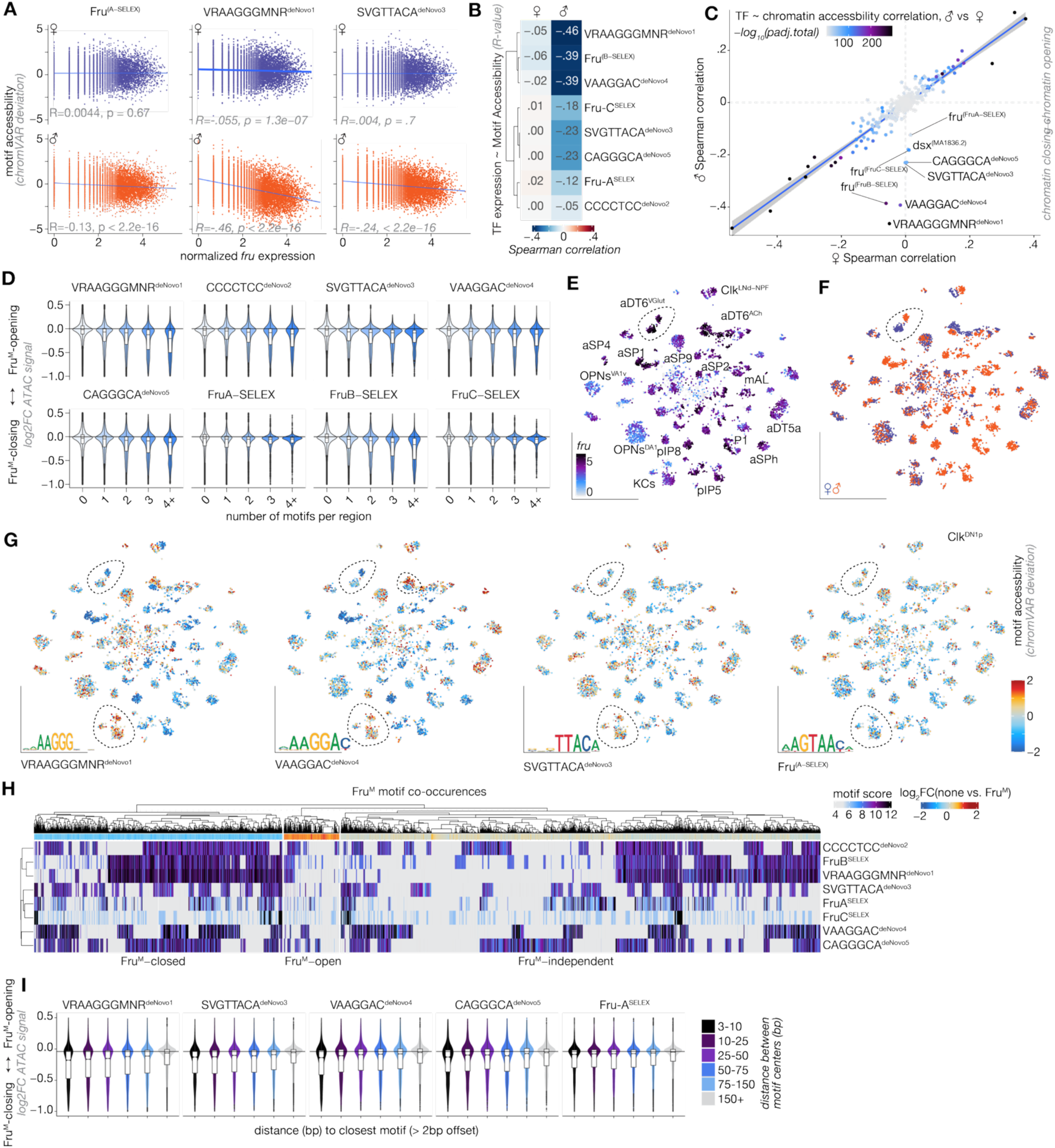
Dose and isoform usage of Fru^M^ protein across unique sequence landscapes. A) *fru* expression vs chromVAR deviations for canonical Fru^M^ motifs across single cells split by sex. B) Heatmap of Spearman R-values across all Fru^M^ motifs, hierarchically clustered. C) Chromatin opening or closing associated R-values across all TF-motif pairs for male cells versus female cells. D) Log2FC of Fru^M^-dependent bulk ATAC chromatin accessibility of regions based on motif identity and multiplicity. All motifs passing FIMO p-val < 1e-3 are counted. E) RNA tSNE of *fru* expression across subset annotated cells. F) RNA tSNE of sex composition across subset annotated cells. G) chromVAR accessibility scores for select Fru^M^ motifs across RNA tSNE subset annotated cells. H) Heatmap of best *de-novo* Fru^M^ motif scores across Fru^M^-closed regions, thresholded by significant motif hits (pval < 0.001). I) Violin plot of bulk ATAC log2FC for Fru^M^ vs. no Fru^M^ neurons stratified by inter-motif offsets.

Because dosage effects for TFs often arise from the ability to bind low-affinity sites at higher doses (Katsani, 1999), we looked at the repressive effect of Fru^M^ on regions with different multiplicities of motifs using our bulk ATAC data. We find a linear effect of increasing the number of binding sites on the strength of closing for each Fru^M^ motif (Figure 6D). Putative regulatory elements rarely have multiple high-affinity matches. Above two motifs, this effect is driven by low-affinity matches (1e-4 >= p-val < 1e-3). In addition to variable dosage as a source of cell-type specific Fru^M^ activity, isoforms of Fru^M^ have been shown to be expressed in different combinations across neuron types (Neville et al., 2014; von Philipsborn et al., 2014). While we can observe RNA reads in the 3’-end DNA binding domain isoform exons of *fru* using 10X technologies, reads in these unique exons are not frequent enough to robustly determine the isoform repertoire expressed in each cell type. As an alternative strategy, we calculated the accessibility of enhancers containing motifs for Fru-A, Fru-B and Fru-C across cell types using chromVAR, as we did previously across all TFs (Figure 3, Figure 6E-G, Supplemental Figure 7C). For example, aDT6 neurons, which are reported to express Fru-B and Fru-C isoforms but not Fru-A (von Philipsborn et al., 2014), show differences in chromatin accessibility between male and female neurons for B (Fru-B^SELEX^, VRAAGGGMNR^deNovo1^) and C (Fru-C^SELEX^, SVGTTACA^deNovo3^) motifs and none for Fru-A motifs.

Finally, we tested for synergistic effects of Fru motif co-occurrence within the same region (annotation synergy), using chromVAR (Supplemental Figure 7B). We find particularly strong synergy between Fru-A and Fru-B motifs, and Fru-B and Fru-C motifs, while the accessibility of sites with these motifs is not highly correlated across cells. This suggests that Fru^M^ action within a cell type is restricted further by the presence of sites with composite motifs which could likely only bind certain isoform heterodimers of Fru^M^. Given the combinatorial expression of Fru isoforms across neuron types, we would expect regulatory regions with composite sites to indeed be far more variable in chromatin accessibility than those without. In our motif analysis of Fru^M^-closed regions in bulk, we find that many sites contain matches for more than one distinct Fru^M^ motif, and the repressive effect of motif combinations is most apparent only when motifs are in close proximity (< 10-25 bp between motif centers) on the DNA (Figure 6H, I). Thus, in addition to the pre-existing cell-type specific chromatin opening landscapes and “closing power” that varies with Fru^M^ dose, cell-type specific gene regulatory activity of Fru^M^ also depends on interaction of different DBD mixes, and all these coalesce to modify enhancers in a way that depends on cis element sequence.

## Discussion

Here, we consider the repressive activity of Fru^M^ in the context of the lineage-fabricated chromatin landscape upon which it acts through the course of development. We find that chromatin effects of Fru^M^ begin to show around 24h APF, are fully established by 48h APF, and durably maintained into the adult. In the adult, these Fru^M^-closed enhancers, opened earlier in development, have highly cell-type specific accessibility in female neurons, suggesting that Fru^M^ closes subsets of lineage-programmed enhancers that give individual neuron types their unique features. Among the set of regulatory elements opened by the cell’s specific history, those that are Fru^M^-closed are those that can win the competition for available Fru^M^ protein dose and DNA binding domain isoforms. Elements that appear to open in the presence of Fru^M^ —i.e. in male *fruitless* cells--are simply hallmarks of cell types absent in the female.

*We thought understanding fruitless neurons would be simpler than the entire cerebrum, but understanding fruitless neurons required understanding the entire cerebrum*

Sex differentiated circuits have been at the forefront of our understanding of neural circuit function for 20 years. The presence of master regulators of sex differentiation advanced studies of instinctual behavior by shining a spotlight on neurons most likely to play causal roles. At the same time, anatomic sex differences within these circuits immediately linked aspects of form and function. We therefore asked how this single transcription factor acts at the molecular level to produce those anatomic and physiological sex differences (Brovkina et al., 2021). For example, we hypothesized that Fru^M^ could cause diverse *fru*-expressing neurons to connect specifically in the male by inducing some kind of universal synaptic glue.

Instead, both Fru^M^ and Fru^COMM^ turn out to be among the strongest transcriptional repressors defined by expression ∼ motif accessibility correlation (Brovkina et al., 2021; Calderon et al., 2022). This means that Fru’s regulatory logic depends fundamentally on its interaction with events produced by activating transcription factors. Because *fruitless* neurons come from diverse stem cell lineages and are precise birth order subtypes within those lineages, understanding how Fru^M^ sex-differentiates gene expression required a cell-type specific point of view that placed it in context of other patterning events. We therefore identified these mechanisms that diversify neurons according to the spatial and temporal origins (Elkahlah et al., 2025). Now, we can see that elements that Fru^M^ can repress were opened in cell-type specific patterns, and we infer that these landscapes are generated by hemilineage-defining transcription factors. Moreover, the genes regulated by Fru^M^ are specific to birth order cohorts within hemilineages. We find here that many birth order factors are also predicted as transcriptional repressors. As some are BTB domain TFs closely related to Fru^M^, one possibility is that birth order TFs and Fru^M^ can close some of the same lineage-patterned enhancers; if a birth order TF does so in a sex-shared manner, there’s nothing left for Fru^M^ to do. Alternatively, Fru^M^ and birth order repressors could work at different sites within the same gene.

Our work raises two major questions: First, how is *fruitless* itself transcriptionally regulated? Both our prior transcriptional analysis and the enhancer landscape of this locus that we report here suggest that each cell type turns *fru* on by different means (Brovkina et al., 2021; Elkahlah et al., 2025). The lack of non-*fru* cells in our dataset limits our ability to make these regulatory links at this specific locus, though we infer that both lineage and birth order TFs must contribute. As we show here, the issue is not simply to turn the locus on or off in such a precise and diverse repertoire of cells, but to regulate transcriptional dose and post-transcriptional splicing of the DNA binding domains.

Second, while adult chromatin appears to be a record of events that occurred over the course of development, we have been surprised to find that the repertoires of transcription factors that reflect hemilineage and birth order identity—and that are predicted to set up the landscape that Fru^M^ acts on—in many cases remain expressed for the lifetime of the neuron (Allen et al., 2026a; Cachero et al., 2025; Elkahlah et al., 2025). Additional transcription factors are modulated by maturation (Elkahlah et al., 2025; Jain et al., 2022). Do Fru^M^ and activating factors do their work at particular enhancers transiently, and then these enhancers are “settled?” Or, do these factors continue to compete at sites they jointly regulate, even into adulthood? Understanding these kinetics will require both developmental chromatin accessibility and timed manipulation experiments.

### Regulatory grammar of Fru^M^ proteins

Different *fru* cell types are known to produce different repertoires of Fru-A, -B, and -C isoforms, and Fru-B and Fru-C are required for courtship behavior, while Fru-A is not (Neville et al., 2014; von Philipsborn et al., 2014). We find that Fru-B sites are sufficient for robust chromatin closing and that closing at Fru-C sites is detectable, while Fru-A sites have modest effects on their own. It seems likely that the stoichiometry between Fru-A, Fru-B, and Fru-C has a significant influence on what sites are being bound, though it remains to be seen if the composition of Fru^M^ dimers are regulated or specific in any way between Fru^M^ isoforms, or if the combinations formed are simply limited by what the cell type produces. *In-vitro*, the BTB-domain of Fru^M^ appears to be less interchangeable with other factors, suggesting that dimerization is far more likely to prefer Fru^M^-Fru^M^-homodimers rather than dimers with other BTB-containing proteins (Bonchuk et al., 2024). Given the strength of Fru-B motifs in attracting chromatin closing, and the synergy of Fru-B motifs with both Fru-A and Fru-C, it is possible that dimers containing at least one copy of Fru-B are most influential. The issue of Fru autoregulation is ever-present.

Studies from other BTB-ZF transcription factors suggest a rather complex regulatory grammar based on the formation of multimers. The BTB-ZF GAF/Trl maintains promoter accessibility in an “anti-repression” mechanism by blocking H1 binding, and it does this through the formation of oligomers (Katsani, 1999; Kerrigan et al., 1991; Omichinski et al., 1997). Non-oligomeric GAF proteins have a strong preference for GAF motifs and bind poorly to variations of the motif, while oligomeric forms of GAF bind strongly to sequences of multiple sub-ideal motifs in tandem (Katsani, 1999). Vertebrate BTB-ZFs have been thought to prefer dimer configurations, but recent work has shown that they too can polymerize – polymerization increases DNA binding and repressive activity at loci with clusters of homotypic motifs in a dose-dependent manner (Mance et al., 2024; Park et al., 2024). This is particularly inspiring in the context of our findings that, across cell types, those with the strongest *fru* expression had the most gene regulatory differences. The motif specificity we find contrasts with more pervasive Fru binding measured by DAM-ID (Neville et al., 2014) and suggests Fru^M^’s regulatory output may be more stringent than DNA binding.

Whether this dose-dependency of Fru^M^ is functional *in vivo* in terms of binding potential, oligomer formation, and repressive capacity and whether this mechanism is developmentally relevant are critical questions for follow up. At the level of behavior, the study of hypomorphic *fru* mutations tells us that during development, only a little amount of Fru^M^ is needed to pattern core courtship behavior: The original *fru* mutations resulted in males that could court but were unable to perform mate discrimination, resulting in characteristic male-male courtship chains (Hall, 1978; Ito et al., 1996). In contrast, null mutants cannot court at all (Demir & Dickson, 2005). Reporters in the *fru* locus, such as the one we use in our analysis here in the heterozygous context, are usually null or hypomorphic alleles. Animals heterozygous for these alleles have circuitry and behavior indiscriminable from wild type, and a halving of *fru* dose pales in comparison to the 20-fold difference in *fru* expression we observe across *fru* cell types. At the cellular level, single-clone labeling in *fru* mAL neurons reveals that while *fru* hypomorphic mutations result in a population of mAL neurons that appear intersex, single-neuron labeling approaches reveal that this is a heterogenous population of neurons with either male-typical or female-typical anatomies (Ito et al., 2012). It appears that the function of Fru^M^ in shaping circuit features has a critical threshold which may be distinct across different neurons and across phases of neuronal maturation.

### Gene expression changes in the context of cell identity and cell death

Lastly, sex determination can induce large changes in gene expression and circuit architecture/function simply by changing programmed cell death, which is the molecular fate of neurons in many lineages (Jiang & Reichert, 2012; Pinto-Teixeira et al., 2016; L. Zhou et al., 1995). In addition to cases where neurons of a particular hemilineage die in one sex or the other, in some cases, females and males appear to retain different subsets of neurons from the same hemilineage (Allen et al., 2026b). This could be due to multiple sex-specific apoptotic events that affect distinct birth order cohorts in the different sexes, or to differential neurogenesis (Ren et al., 2016).

Neurons differentiated by Fru^M^ are also differentiated across axes of birth order and lineage (Allen et al., 2026b; Cachero et al., 2025; Elkahlah et al., 2025). Because *fru* expression turns on relatively late in neuronal maturation, it likely arrives on the scene after lineage and birth order factors have generated cell-type specific chromatin landscapes; however, as we describe above, many of these factors, once induced, continue to be produced for the life of the neuron. We expect that lineage-programmed apoptosis opens enhancers in pro-apoptotic genes, and that one of Fru^M^’s most important functions is to close these enhancers or prevent them from opening; as we show, shielding these cells from apoptosis appears to require a particularly high *fru* dose. Because the cells that would have activated these enhancers die, we have not been able to map these enhancers in our current dataset. While lineage-based strategies for blocking apoptosis in neurons and sequencing the populations is feasible (Elkahlah et al., 2025; Ren et al., 2016), this strategy does not work under the control of *fru^P1^*-GAL4, which likely activates p35 too late (data not shown).

In addition, we find many cases in which Fru^M^ appears to regulate enhancers in birth order transcription factor loci. Adjusting the repertoire of fates within hemilineages could be another way for Fru^M^ to produce large sex differences in neurons or circuits through small primary adjustments of gene expression. However, we do not find a rule for this across cell types nor a single temporal window that *fru* occupies (Figure 5) (Elkahlah et al., 2025). We caution that in bulk RNAseq analyses or in single-cell analyses that don’t match birth order cohorts within hemilineages, differences in cell type distribution due to apoptosis or other mechanisms overwhelm true, within-cohort sex differences. Parsing direct targets versus indirect downstream consequences is crucial for understanding transcriptional mechanisms through which master regulators of sex determination influence development.

## Acknowledgements

We thank members of the Clowney lab for participating in bulk brain dissections for sorting experiments; Abbigayl Burtis for assistance in preparation of bulk ATAC- and RNA-seq libraries; and Yijie Pan for generating connectome cartoons. FAC-sorting was performed with Kamlai Saiya-Cork in the BRCF Flow Cytometry Core. Library prep and next-generation sequencing were carried out in the Advanced Genomics Core at the University of Michigan. Research reported in this publication was supported by the University of Michigan Advanced Genomics Core, the UM Single Cell Spatial Analysis Program and the National Cancer Institutes of Health under Award Number P30CA046592 using the following Cancer Center Shared Resource: Single Cell and Spatial Analysis Shared Resource. Jessica Tollkuhn, Kevin Monahan, Yerbol Kurmangaliyev, and Alexander Marande provided technical input on analyses and/or feedback on the manuscript. Funding was provided by NINDS Ruth L. Kirschstein NRSA 1F31NS127484, Cell and Molecular Biology Graduate Program NIH T32 GM145470, and Hearing Balance and Chemical Senses NIH T32 DC000011 to MVB; by McKnight, Pew, and Rita Allen Scholar awards to EJC; and by NIH R01DC018032 and the University of Michigan.

## Methods

**Table 1:**
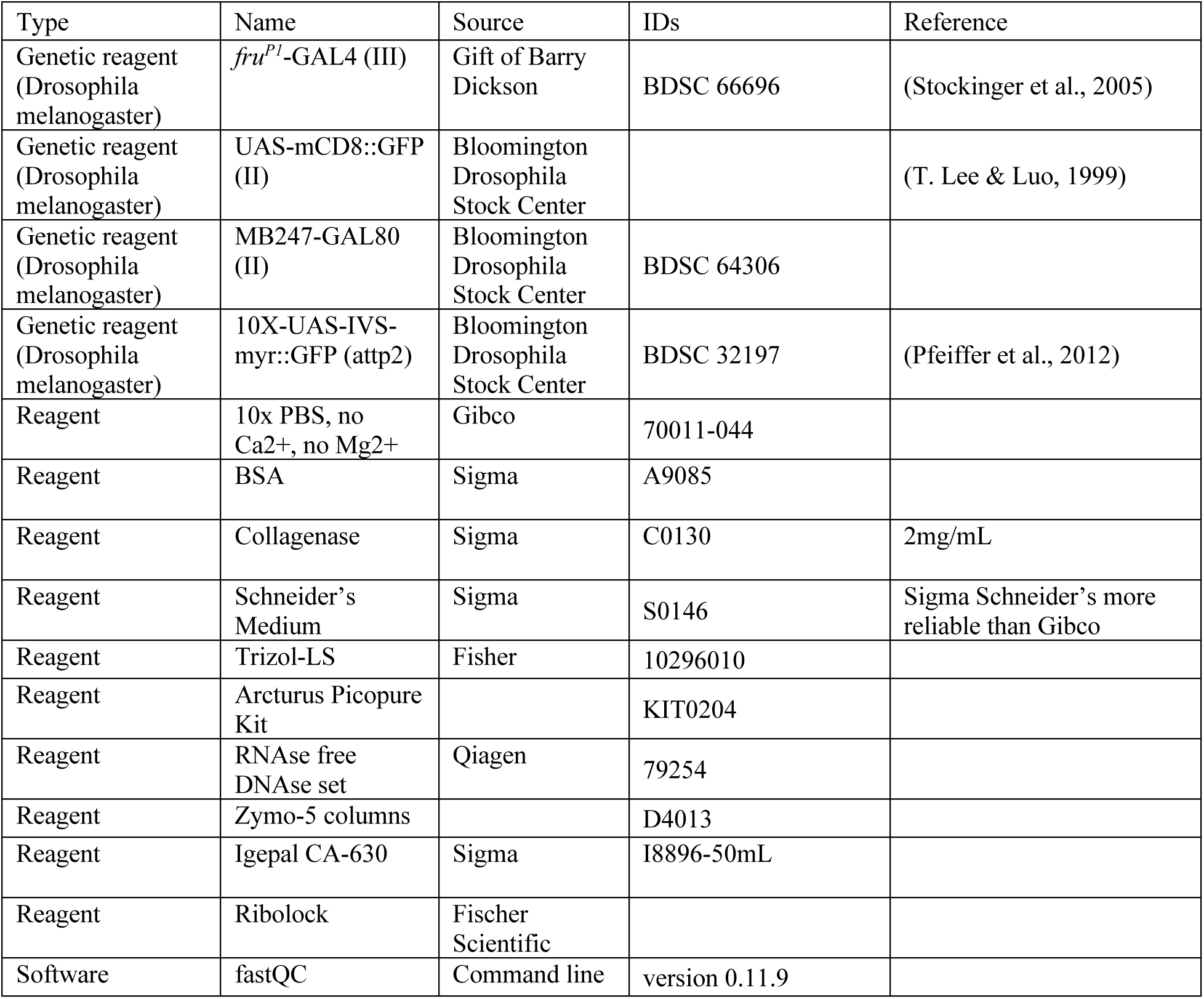
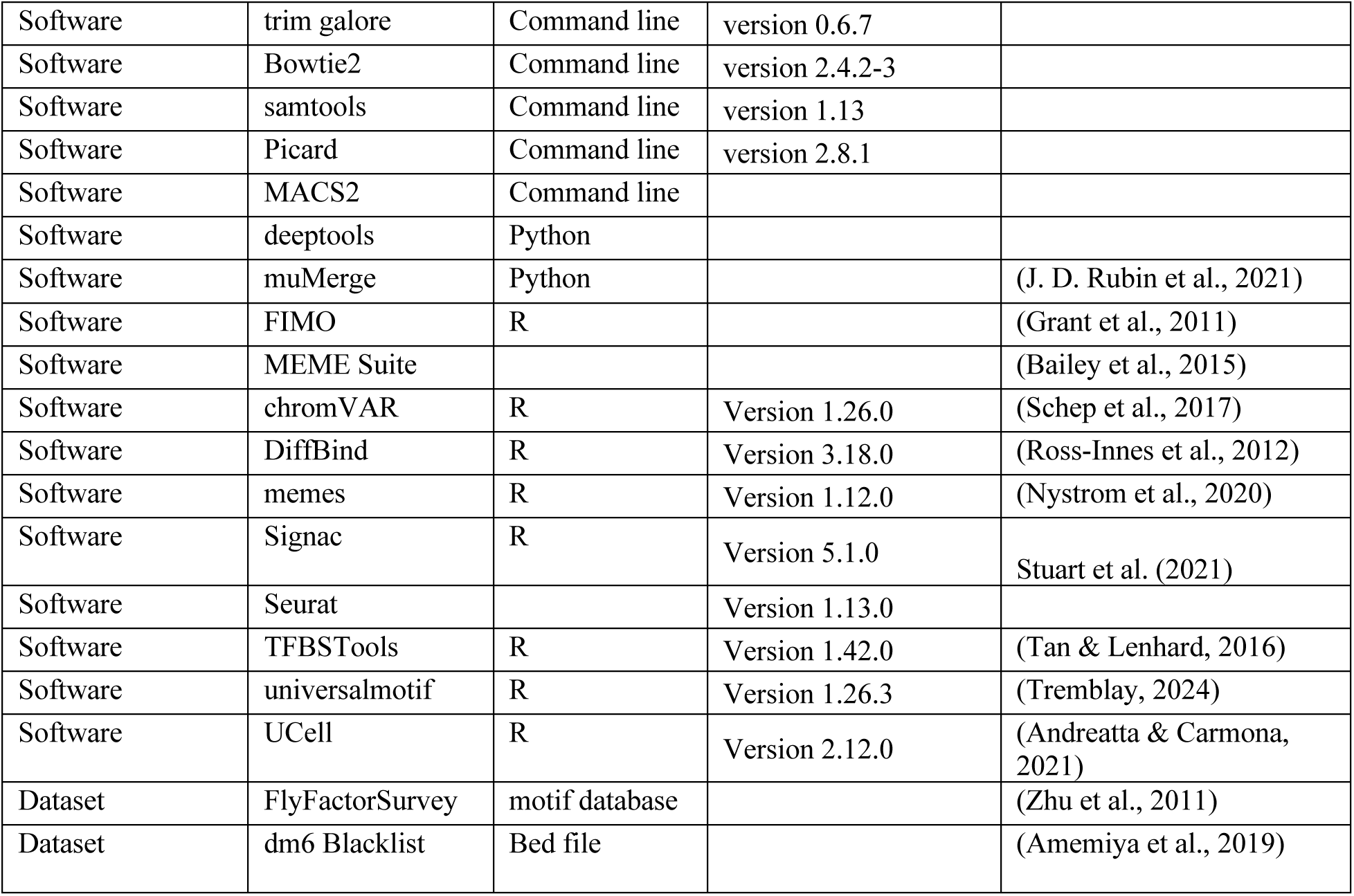
Reagents used.

### Connectome Images

We used a custom python script to download EM skeletons associated with annotated fru-expressing neurons and the rest of their associated lineages from the FAFB v783 (Deutsch et al., 2025; Dorkenwald et al., 2024; Matsliah et al., 2023; Schlegel et al., 2024) and MCNS 0.9 (Berg et al., 2025) connectomes, register them onto JFRC2018U, and plot them using navis (Schlegel et al., 2023).

### Flies

Flies were maintained on cornmeal food with yeast sprinkles (‘B’ recipe, Lab Express, Ann Arbor, MI) in a humidified incubator at 25C on a 12:12 light:dark cycle. Adult flies were collected as 2-7 day old adults who were housed in mixed-sex groups. 48h APF and 24h APF animals were collected as late wandering L3 larvae, split by sex based on visual presence of larval gonads, then allowed to mature for 48h or 24h, respectively in a humidified incubator at 25C on a 12:12 light:dark cycle.

**Table 2:**
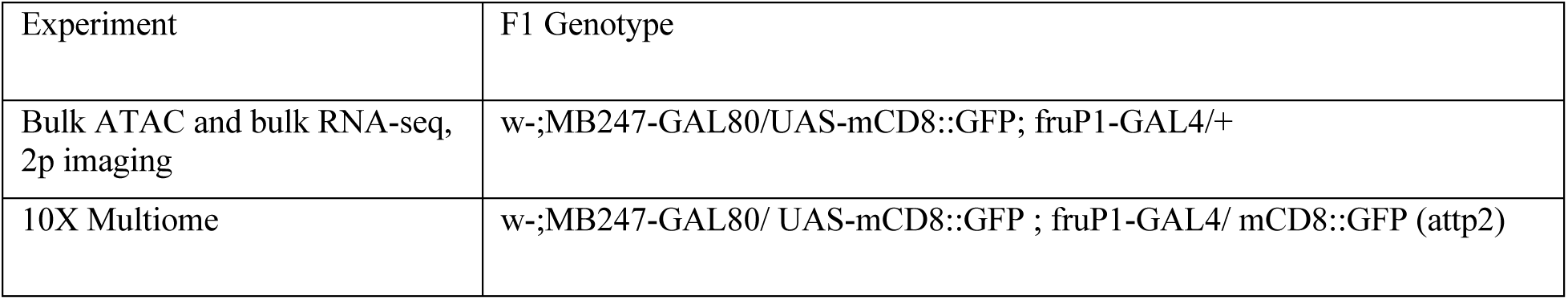
List of Genotypes.

### Brain dissections for imaging

Brains were dissected in external saline (108 mM NaCl, 5 mM KCl, 2 mM CaCl2, 8.2 mM MgCl2, 4 mM NaHCO3, 1 mM NaH2PO4, 5 mM trehalose, 10 mM sucrose, 5 mM HEPES pH7.5, osmolarity adjusted to 265 mOsm). For two-photon imaging, brains were then transferred fresh to 35mm imaging dishes and pinned to sylgard squares with tungsten wire. Imaging was performed on a Bruker Investigator using a 1.0 NA, 20x, water-dipping objective. Stacks were collected along the anterior-posterior axis with 1 micrometer spacing in Z and ∼350nm axial pixel size.

### Flow Cytometry

Brains were dissected for up to 90 minutes in Schneider’s medium supplemented with 1% BSA and placed on ice. Optic lobes were removed during dissection. Brain dissections were interspersed such that both male and female brains were dissected throughout the dissection period. For pupal timepoints, 30-60 brains were obtained for each sex. For adult 10X Multiome, 60-100 mixed sex animals were obtained per replicate. After all dissections were completed, collagenase was added to a final concentration of 2mg/mL and samples were incubated at 37C for 12 minutes (pupal brains) or 20 minutes (adult brains), without agitation. Samples were dissociated by trituration and spun down at 300g, 4C, for 5 minutes. Collagenase solution was removed and replaced with PBS+0.1% BSA, and cells were passed through a cell strainer cap and supplemented with 50ng/mL DAPI immediately before being subjected to flow cytometry. Cells were sorted at the UMich Flow Cytometry Core on a FACS Aria II for replicates 1 and 2 (8344-MB and 8506-MB) and on a BD FACS Discover 8 for replicates 3-5 (10690-MB, 10971-MB, 11042-MB) due to the FACS Aria II being decommissioned. Plasticware for cell dissociation and collection was pre-treated by rinsing with PBS+1% BSA to prevent cells from sticking to bare plastic.

During flow cytometry, dead and dying cells were excluded using DAPI signal, and forward scatter and side scatter measurements were used to gate single cells. Using our dissociation methods, 80-95% of singlets appeared viable (DAPI-low), and 2-5% of viable singlets were GFP+. We collected 2,000-5,000 GFP+ cells for each *fru* sample and analyzed 10,000 GFP-negative (*fru*-negative) cells. For each replicate, we sorted male and female cells during the same session and performed transposition or RNA extraction in parallel. To protect the dissociated fly primary cells on the FACS Aria II, we disabled agitation of the sample tube, and sorted using the “large nozzle,” e.g. 100 *μ* m, i.e. using larger droplet size and lower pressure. For ATAC-seq and 10X Multiome, we sorted cells into PBS supplemented with 0.1% BSA. For RNA-seq, we sorted cells directly into Trizol-LS.

### Bulk Assay for Transposase-Accessible Chromatin (ATAC)

After FAC-sorting, nuclear isolation and Tn5 transposition were performed as in (Buenrostro et al., 2015) with modifications made for small numbers of cells and for *Drosophila* (Brovkina et al., 2021; Merrill et al., 2022; Monahan et al., 2017). Transposed DNA was isolated and stored at −20C until library preparation. Transposed DNA was amplified with barcoded primers in NEBNext High Fidelity 2X PCR Master Mix (NEB) and purified with Ampure XP beads (Beckman Coulter) at a ratio of 1.6uL beads per 1uL library. Purified library was eluted in 30uL of 10mM Tris-HCl pH8, 0.1mM EDTA. The quality of prepared libraries was verified with a Bioanalyzer 2100 using a high sensitivity DNA kit (Agilent) at the University of Michigan Advanced Genomics Core. Libraries were quantified using KAPA qPCR assay (KAPA Biosystems), multiplexed, and sequenced on a NextSeq Illumina machine with 75 base pair, paired-end reads. Libraries were sequenced to a depth of 35-70 million reads.

### ATAC-Seq Processing

Quality control was done using fastqc (version 0.11.9). Adapters were trimmed using trimgalore (version 0.6.7) and reads below 20 bases were discarded. Reads were aligned to dm6 with Bowtie2 (version 2.4.2-3) with the following parameters: option –X 1000 to set the maximum fragment length for paired-end reads, -p for paired-end alignment, and --no-unal to suppress writing unaligned reads. Aligned reads were processed with samtools (version 1.13) to create a bam file with indexing. Blacklist regions for dm6 were masked (Amemiya et al., 2019). Picard MarkDuplicates (version 2.8.1) was used to mark duplicates and samtools (version 1.13) was used to generate a final bam file with de-duplicated reads, passing a q30 quality filter, aligned to standard chromosomes (chr2L chr2R chr3L chr3R chr4 chrX chrY), and with fragments of 150bp or less to exclude nucleosomal reads. After stringent filtering, we obtained 10-25 million unique reads per library. For downstream differential binding analysis, bam files were also split into X chromosome files and autosome files.

To call peaks, filtered bam files were converted to single-end bed using bamtobed, and peaks were called with MACS2 “callpeak” using –nomodel at an FDR threshold of <0.001, which gave reproducible and robust peak calls across replicates and by eye (Feng et al., 2011). Coverage tracks were generated on filtered bams using deepTools bamCoverage with a window of 1. All ATAC-seq processing was performed on the Great Lakes High Performance Computing Cluster. A consensus peakset was generated using muMerge of all peaks (shared and singletons) called on pooled samples.

### Analysis of Differential Chromatin Accessibility

X chromosome reads and autosomal reads were processed independently using Diffbind in R, as we described in Brovkina et al., 2021. Peaks across samples were turned into a common peakset of regions present in more than one sample and extended to 200bp flanking a calculated consensus summit by default parameters. The consensus peakset generated by DiffBind was exported and used further for feature matrix generation in the 10X Multiome data. Counts from autosomal and X-chromosome split bam files were quantified separately across the 40,091 consensus sites and normalized for read depth. Custom contrasts were added to dba.analyze to compare across neuron type, *fru* status, sex, and developmental time. We primarily analyzed Fru^M^ neurons vs those without Fru^M^ across all timepoints. This is distinct from our previous analysis in the adult, where we took the intersection of chromatin accessibility changes between male *fru* vs female *fru* and male *fru* vs male non-*fru* (Brovkina et al., 2021). A cutoff threshold of FDR < 0.01 was used as the significance cutoff for all contrasts. Unfiltered DiffBind comparisons from X chromosome and autosome analysis were concatenated (Supplemental Table 1). The consensus peakset generated by DiffBind was exported and used further for feature matrix generation in the 10X Multiome data.

Regions from DiffBind were annotated using ChipSeeker based on the nearest gene (G. Yu et al., 2015). Nearest genes were collapsed into lists of unique genes for GO analysis using DAVID. For GO analysis, genes with Fru^M^-open peaks and genes with Fru^M^-closed peaks were compared to a background of genes annotated across all peaks. Normalized counts exported from DiffBind were averaged across replicates of the same samples. To generate z-score heatmaps, mean counts tables were scaled across samples per site, allowing the chromatin opening and closing dynamics of individual genomic regions to be compared across samples.

### Motif analysis

We used the memes package in R to interface with the MEME suite database (Bailey et al., 2015; Nystrom et al., 2020). A custom motif file was built from combined motifs from FlyFactorSurvey, cisBP, and transfac (Zhu et al., 2011) with specific addition of the FruA, FruB, and FruC position-weighted matrices from SELEX analysis (Dalton et al., 2013), as well as one for Prospero (Hassan et al., 1997). We searched for 7-13bp motifs enriched in Fru^M^-open or Fru^M^ closed regions compared to Fru^M^-independent regions using STREME in R (Bailey, 2021; Nystrom et al., 2020). We did not rebase or center peaks for motif analysis, as recommend for ATAC-seq. *De novo* motifs were searched against the motif database using TOMTOM. To identify the positions and significance of each *de novo* motif across all our open chromatin regions, we ran FIMO (Grant et al., 2011) with a p-value threshold of 0.001 (Supplemental Table 2). Hits passing 1e-3 were “weak” matches, while hits passing 1e-4 were “strong” matches. Both *de novo* and SELEX Fru motifs were correlated in PWM space using universalMotif CompareMotifs PCC correlation. To determine distance relationships between motifs, we restricted analysis to motif matches within a peak and calculated the distances between the center of each motif to all other motifs within a peak.

### Bulk RNA Sequencing

Cells were sorted into Trizol-LS and stored at −80C until RNA isolation. For RNA extraction, we followed the standard Trizol-LS protocol until the aqueous phase was isolated. We then passed the aqueous phase over Arcturus Picopure columns, including a DNAse treatment on the column as previously described (Brovkina et al., 2021). After RNA isolation, samples were inspected for quality on a TapeStation. Library preparation was performed using the NEBNext Single Cell/Low Input RNA Library Prep Kit for Illumina, with amplification cycles calibrated to the amount of total RNA. Unstranded, poly-A selected libraries were sequenced on an Illumina NovaSeq using 150bp paired end reads to a depth of ∼30 million reads per library.

After sequencing, all processing steps were done through the Galaxy web platform (The Galaxy Community et al., 2024). Raw reads were trimmed using trim galore! with automatic adapter detection and aligned to dm6 using HiSAT2 in paired-end mode. Uniquely aligning reads above MAPQ 30 were selected using JSON and MarkDuplicates, retaining 20-30 million reads per library. FastQC and MultiQC were used to visualize quality metrics across samples. Coverage tracks were generated using deeptools “bamCoverage”. Exon reads were counted and aggregated at a gene-level using featureCounts. TPM values were calculated using StringTie and averaged between replicates and normalized counts were used for downstream analysis (Supplemental Table 9). Differential expression was determined using DESeq2 using an p.adj threshold of < 0.05 for reporting significance (Love et al., 2014).

### 10X Multiome Sequencing

Adult flies 2-7 days of age of the genotype UAS-mCD8::GFP, MB247-Gal80; *fru^P1^*-GAL4, UAS-mCD8::GFP (attp2) were dissected and sorted on GFP expression as described above. Male and female brains were dissected separately. In our first three libraries, they were then pooled before sorting to prevent batch effects and male-female comparisons being confounded. For the 4^th^ and 5^th^ libraries, we kept the two sexes separate. A total of 20,000–80,000 GFP+ cells were collected into 300uL of 1X PBS + 0.1% BSA. Samples were taken to the Advanced Genomics Core and whole cells were spun in a benchtop Eppendorf centrifuge for 800g x 8 minutes at 4C, similar to bulk-ATAC protocol performed in the fly optic lobe (Jain et al., 2022). After supernatant was carefully removed, cells were resuspended in 10ul 10x Genomics dilute nuclei buffer supplemented with 1U/uL of Ribolock RNase inhibitor (Thermofisher Scientific, catalog no. FEREO0382) and quantified on a Luna cell counter. 10K cells were targeted with an average of 50% overload; we recover a range of 3-7k cells per library.

Live cells were input directly into the transposition reaction with no nuclear isolation or detergent permeabilization and prepared using the 10x Genomics Single Cell Multiome ATAC + Gene Expression kit (1000285). We presume that the membranes of *D. mel* cells, kept at 25C normally and then kept on ice for the entire flow cytometry and transport processes, become permeable at 37C. This protocol allowed us to recover a reasonable number of sorted cells and improved RNA recovery dramatically, as fly nuclei have little RNA. We refer to our data as scRNAseq/ATACseq, rather than snRNAseq/ATACseq.

Library prep and next-generation sequencing was carried out in the Advanced Genomics Core at the University of Michigan. The GEX pool was subjected to 151×151bp paired-end sequencing, while the ATAC pool was subjected to 51×51bp paired-end sequencing, according to the manufacturer’s protocol (Illumina NovaSeq) to a target read depth of 50k reads per cell for both modalities. BCL Convert Conversion Software v4 (Illumina) was used to generate de-multiplexed Fastq files. ATAC and GEX fastq files were processed using CellRanger ARC Analysis using noncoding RNAs and intronic reads against a custom dm6 genome reference with scaffolds for reporter alleles GAL4 and mCD8::GFP.

### Multiomic Analysis – Quality Control

Initial matrices were generated with unfiltered data with at least 300 ATAC fragments and 200 detected genes. ATAC matrices were generated in two ways: First, using 2kb genomic bins with blacklisted regions masked. This analysis was used for weighted-nearest neighbor analysis and clustering to maximize information from rare cell types that would be missed using a peakset. Second, we generated counts matrices across the consensus peakset output from our bulk ATAC-seq, which were generated in directly comparable cells at much higher coverage and depth. RNA data was added for each library and libraries were merged into one object.

Annotations were made using an Ensembl annotation hub made from *Drosophila_melanogaster.BDGP6.32.104* (AH92037) in NCBI format. We obtain distinct TSS enrichment, though systematically lower than mammalian cells. We also see characteristic nucleosomal periodicity in fragment size indicative of an appropriately transposed ATAC-seq experiment. We obtain a median of 2000 unique gene features per cell, which is on par with whole-cell scRNA-seq done in *fru* neurons in pupal stages, though we do get more mitochondrial genes than typical (Palmateer et al., 2023). This is unsurprising, given that our cells are cooked at 37C for 30 minutes during bulk ATAC transposition. Given the quality tradeoff in nuclear RNA coverage, quality, and simple biological relevance in traditional multiomic data, the quality difference was acceptable. Thus, scATAC-seq on whole cells appears to work as well as snATAC-seq on isolated nuclei in *Drosophila*, likely due to heat-shock of the membranes and small cell size compared to mammalian cells for which the platform is optimized.

Because we used an expanded transcript reference compared to other Drosophila melanogaster single-cell RNA datasets, we kept protein-coding genes and long non-coding RNAs in the gene expression matrices, filtering out transposable elements, mitochondrial genes, small RNAs, pre-RNAs, pseudogenes, and Stellate elements on the Y chromosome.

Cells were filtered on the following metrics: nCount_bins < 25000 & nCount_RNA < 55000 & ATAC_counts_per_feature < 1.75 & Xratio < 0.28 & Xratio > 0.1 & TSS.enrichment < 4 & nucleosome_signal < 1 & nucleosome_signal > .25 & nCount_ATAC >= 250 & nFeature_RNA >= 200. After an initial dimensionality reduction (before any subsetting) and clustering (resolution = 2) according to the section below, we filtered out clusters X0, X1, X4 for clustering poorly and low quality. Dimensionality reduction was rerun on fully subset data. For additional quality control metrics and variables to regress, UCell scores were calculate for transcripts involved in heat-shock (^Hsp), oxidative phosphorylation (^ATPsyn), and neuronal activity genes often upregulated during sample handling (Hr38, sr, CG14186).

### Multiome analysis – Assigning donor sex

To demultiplex male and female cells computationally, we generated a probabilistic score for male sex determination genes *lncRNA:roX1* and *lncRNA:roX2* using UCell and used a cutoff of >= 0.40 to call male cells, calibrated against our male-only and female-only libraries. The sum of ATAC reads from the X chromosome per cell were divided by the sum of autosomal reads to generate an X:Auto ratio. Our UCell calls for male and female showed very distinct distributions of X:A ratios corresponding to the strict bi-modality in X:A ratio split at 0.18. A handful of cells in the female and male-only libraries were manually reassigned. After cells were assigned by sex, ratios of male:female cells per cluster were inspected for correspondence to the literature.

### Multiomic Analysis – Dimensionality Reduction

We performed a peak-independent clustering of ATAC using 2kb genomic bins (Figure 2B). Sex chromosomes were excluded during normalization and clustering to avoid “bi-lobed” clusters driven by differing X:Autosome ratios in *D. mel* (Cusanovich et al., 2018). For peak and bin ATAC analyses, TfIDF, FindTopFeatures and RunSVD were run with 200 dimensions. Batch correction for ATAC was done using anchor integration and Harmony for RNA, after which corrected lsi embeddings (2:100 dimensions) and corrected PCA values (1:100 dimensions) were integrated using a weighted-nearest neighbors approach and clustered. A resolution of 2 generated 224 initial clusters. For RNA dimensionality reduction, we used LogNormalize and ScaleData with the following variables regressed: nCount_RNA, library, percent_mito, percent_ribo, oxp_UCell, hsp_UCell, and arg_UCell. 200 principal components were calculated, and tSNE and UMAP reductions run with 1:100 dimensions. SCTransform was not used due to significant dropout of lower-expression genes, strongly impacting clustering.

We expected ∼60 major anatomically defined types of *fru* neurons in the central brain from the literature. In our clustering, we identify a number of these cell types as large, distinct clusters. Kenyon cells (*ey+, toy+, dac+)* split into three clusters based on birth-order subtypes. 2 types of *fru* olfactory projection neurons (Oaz+, otp+), DA1 (acj6+) and VA1v (*Pax*, *mirr, vvl*) split into two distinct clusters based on known markers (Li et al., 2018). Other cell types like the GABAergic inhibitory gustatory “mAL” interneurons (*Fer1+*, *TfAP-2+, Gad1+)* split into several cluster based on birth-order factors (Elkahlah et al., 2025; Sengupta et al., 2022; Yuan et al., 2013).

We used these cell types to calibrate clustering resolution. The majority of clusters correspond to very small Imp+ clusters, suggesting these clusters correspond to very rare or singleton *fru* neurons not extensively characterized anatomically or transcriptionally; these may be enriched for early-born cells (primary neurons) and may be more likely to be female-specific, while larger and later-born groups of *fru* neurons are more likely to be male-specific (Allen et al., 2026b). Other cell types were determined using markers from the literature and information from the connectome about predicted neurotransmitter identities and hemilineage notations (Supplemental Table 3). For cell types where neurons split into multiple clusters based on birth order, we generate a separate metadata assignment of cell type where subtypes are merged. We were able to identify optic lobe neurons in two ways: label transfer from the 48h *fru* neuron single-cell RNA-seq dataset (Palmateer et al., 2023), where optic lobe types are very large, distinct clusters; and by inspecting library stoichiometry, since optic lobe types only come from our first three independent, mixed-sex replicates. To generate plots of “annotated” *fru* neuron types, we subset out all unnamed clusters, as well as unnamed optic lobe types label-transferred from the 48h APF single-cell *fru* dataset. For the annotated subset, UMAP and tSNE reductions were then re-run using 1:100 dimensions on the RNA assay alone.

### Cell-type specific peaks

To generate cell-type specific ATAC peaks, FindMarkers() was run on clusters at the cell-type level (Supplemental Table 5). The top (by p.adj value) 1k enriched peaks were taken from annotated clusters and condensed into a set of 12.882 unique regions. Annotated cells were pseudobulked by replicate and used to run DoHeatmap() with the 12k regions. Since there is no native way to assign each region to an annotated cell type based on the heatmap, we took the data produced by DoHeatmap and generated a binary matrix of the signal. We ran a custom R function to find binary runs within the ordered matrix and assign them to each cell type.

### Differential gene expression

Differential expression between male and female cells was calculated using pseudobulked data using library replicates. First, cells were aggregated by sex, cluster, and library replicate. Conditions with fewer than 4 cells making up the pseudobulk data were filtered out. Only clusters with at least three library replicates in both male and female conditions were compared using FindMarkers() with the DESeq2 option (p.adj < 0.05) (Supplemental Table 8). Cluster differential expression was done on the “subtype” (i.e., distinct birth order subset within hemilineage) level to preserve comparisons between molecularly matched “type” as much as possible and avoid differential expression being dominated by cases where female neurons and male neurons of a cell type comprise distinct subtypes, e.g. female mAL neurons being solely *br+* due to apoptosis of the *pdm3+* type (Elkahlah et al., 2025). After DESeq2 testing, we merged the differential genes found across different subtypes when possible and for calculations of number of differentially expressed genes.

### Single-cell motif analysis

For motif analysis, ChromVAR was run using the native Signac function against the custom motif database we generated for bulk ATAC-seq analysis, with *de-novo* Fru^M^ motifs added. To calculate motif activity per transcription factor, we ran spearman correlations between the RNA normalized expression of a transcription factor and its motifs across all cells, as well as female and male cells separated (Supplemental Table 6). We ran PANDO (Fleck et al., 2023) with (exons == F) using our consensus peakset and custom motif database. Since Pando aggregates all motifs per gene, our CCCCTCC^deNovo2^ motif was excluded due to its repetitive nature, prevalence, and lack of strong relationship to *fru* expression. To maintain computational power and biological relevance, counts for *fru* expression in female cells were zeroed to reflect lack of Fru^M^ protein activity, rather than filtering on male cells only.

### Computational Analysis of Public Datasets

Table of SRA accessions for previously published ATAC data

| SRA Accession | Sample Name | Reference |
| --- | --- | --- |
| SRR11818939 | ATAC_L3_Neuroblast_II_R1 | Larson et al. 2021 |
| SRR11818940 | ATAC_L3_Neuroblast_II_R2 | Larson et al. 2021 |
| SRR15461247 | ATAC_Embryo_Stg12_Neuroblast_I_R4 | Lucas et al. 2021 |
| SRR15461248 | ATAC_Embryo_Stg12_Neuroblast_I_R3 | Lucas et al. 2021 |
| SRR15461249 | ATAC_Embryo_Stg12_Neuroblast_I_R2 | Lucas et al. 2021 |
| SRR15461250 | ATAC_Embryo_Stg12_Neuroblast_I_R1 | Lucas et al. 2021 |
| SRR15461251 | ATAC_Embryo_Stg10_Neuroblast_I_R5 | Lucas et al. 2021 |
| SRR15461252 | ATAC_Embryo_Stg10_Neuroblast_I_R4 | Lucas et al. 2021 |
| SRR15461253 | ATAC_Embryo_Stg10_Neuroblast_I_R3 | Lucas et al. 2021 |
| SRR15461254 | ATAC_Embryo_Stg10_Neuroblast_I_R2 | Lucas et al. 2021 |
| SRR15461255 | ATAC_Embryo_Stg10_Neuroblast_I_R1 | Lucas et al. 2021 |
| SRR15461256 | ATAC_Embryo_Stg5_Neuroblast_I_R1 | Lucas et al. 2021 |
| SRR18192452 | ATAC_Adult_Neurons_GABA_R3 | Merrill et al. 2022 |
| SRR18192453 | ATAC_Adult_Neurons_GABA_R2 | Merrill et al. 2022 |
| SRR18192454 | ATAC_Adult_Neurons_GABA_R1 | Merrill et al. 2022 |
| SRR18192455 | ATAC_Adult_Neurons_TH_R2 | Merrill et al. 2022 |
| SRR18192456 | ATAC_Adult_Neurons_TH_R1 | Merrill et al. 2022 |
| SRR12248072 | ATAC_L3_Neuron_Female_R1 | Chen D et al. 2021 |
| SRR12248073 | ATAC_L3_Neuron_Female_R2 | Chen D et al. 2021 |
| SRR12248074 | ATAC_L3_Neuron_Female_R3 | Chen D et al. 2021 |
| SRR12248078 | ATAC_82hAPF_Neuron_Female_R1 | Chen D et al. 2021 |
| SRR12248079 | ATAC_82hAPF_Neuron_Female_R2 | Chen D et al. 2021 |
| SRR12248080 | ATAC_82hAPF_Neuron_Female_R3 | Chen D et al. 2021 |

For reanalysis of ATAC-seq data from sorted neuroblasts and neurons, raw fastq files were downloaded with SRA toolkit with the accessions in the table above. For Type II Neuroblast ATAC-seq data, dissected brains of brat*^11/Df(2L)Exel8040^* larvae where most of the brain is composed of Type II NBs were profiled (Larson et al., 2021). Embryonic neuroblast ATAC-seq data was derived from staged embryos FAC-sorted for Deadpan-eGFP (Lucas et al., 2021). Adult neurons were sorted by Gad1-GAL4 for GABAergic neurons and by TH-GAL4 for dopaminergic neurons (Merrill et al., 2022). For L3 and 82h APF female ATAC profiles, neurons were labeled by ELAV-GAL4 and FAC-sorted (D. Chen et al., 2021). All datasets were processed with the same pipeline as the bulk ATAC-seq data generated in this paper.

---

**Supplemental Figure 1.**
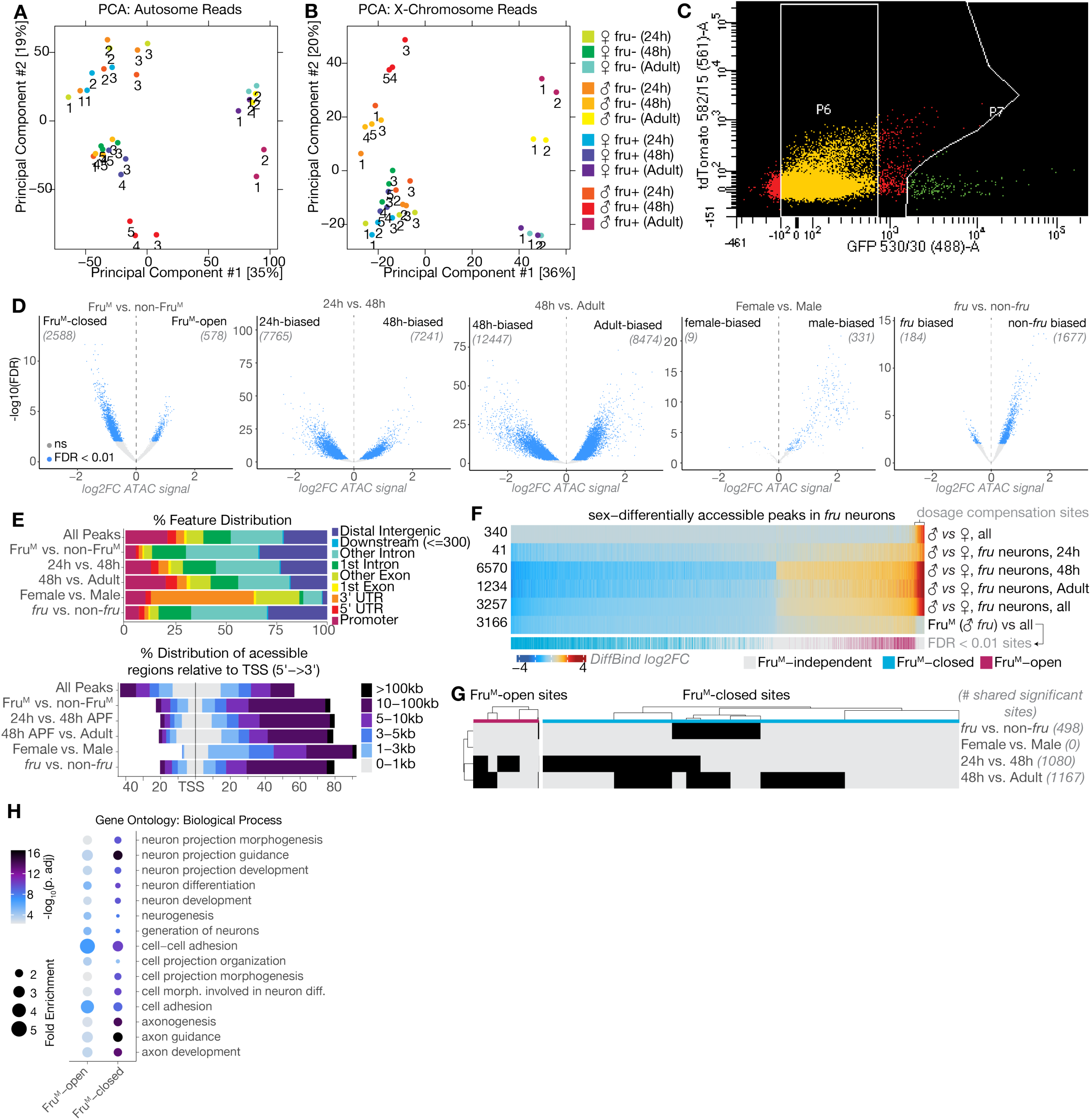
A) Autosome and B) X-chromosome Diffbind PCA plots for all replicates analyzed C) Example FACS plot of sorted *fru*-neurons showing *fru*-negative P6 and *fru*-positive P7 populations. D) Volcano plots of all major comparisons made using DiffBind. E) Annotation of differential peak distribution within genomic locations (top) and as distances from the TSS (bottom). F) Heatmap of log2FC differential accessibility across a union set of significantly changed peaks for comparisons in Fig 1E. G) Binary heatmap of Fru^M^-DARs changing in additional comparisons with number of overlapping sites labeled. H) GO analysis of biological processes enriched in genes with Fru^M^-open or Fru^M^-closed peaks over the set of all genes with accessible chromatin.

**Supplemental Figure 2, related to Figure 1.**
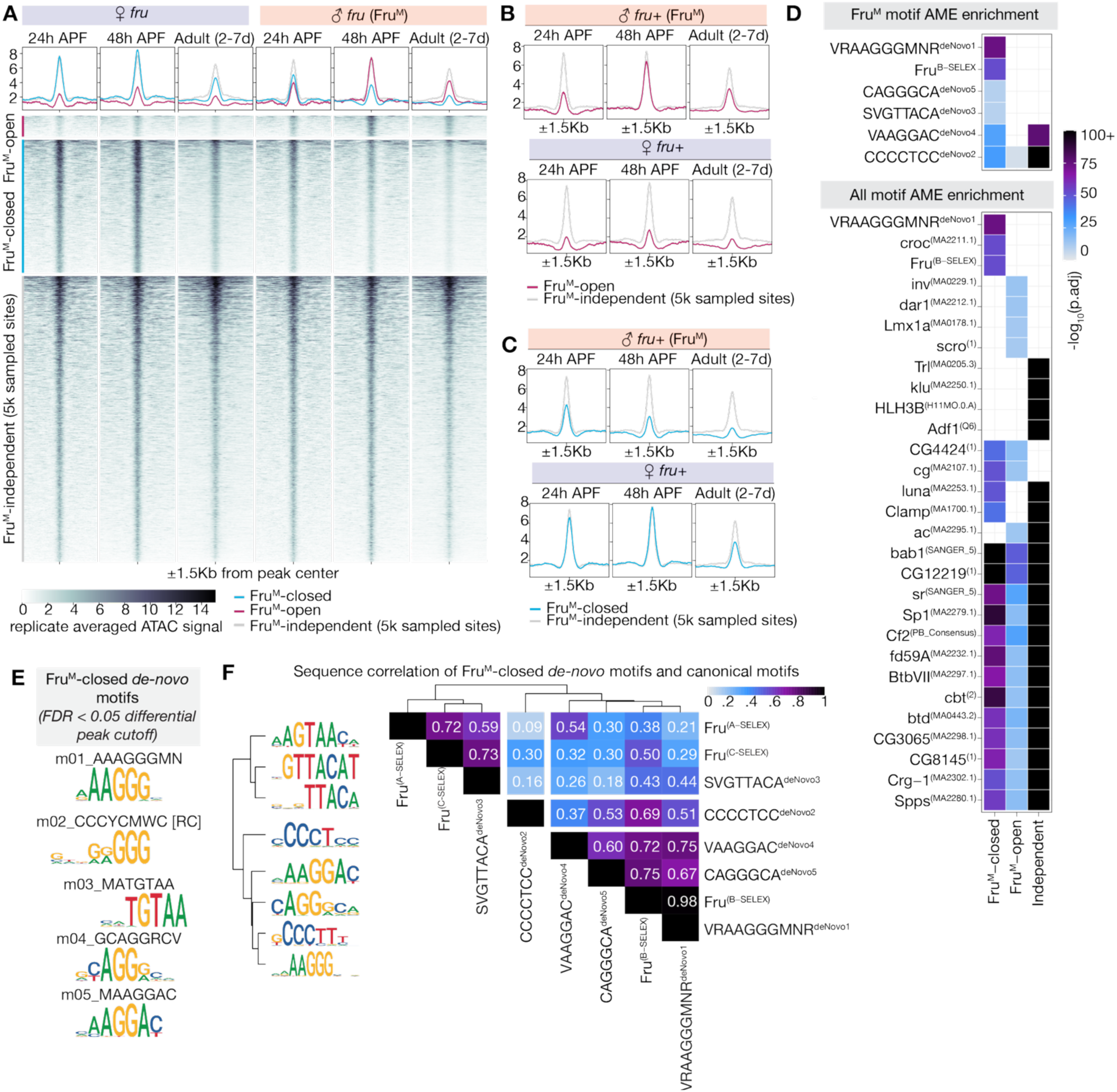
A) DeepTools signal heatmap of replicate-average ATAC signal from Fru^M^-DARs and a 5K random sample of independent regions. B-C) Mean signal from (A) across Fru^M^-open (B) and Fru^M^-closed (C) compared to sampled independent regions. D) Heatmaps showing significance of enrichment using AME with a shuffled background for Fru^M^ *de novo* and canonical motifs (top) and top 15 motifs across all PWMs (bottom). Enrichments are calculated independently for each peak set versus shuffled background E) STREME *de novo* motifs identified with same parameters as Fig 1I at an FDR < 0.05 differential peak threshold. F) PCC correlation showing motif similarity across *de novo* motifs in Fru^M^-closed peaks and known Fru SELEX motifs for A, B, and C isoforms.

**Supplemental Figure 3, related to Figure 2.**
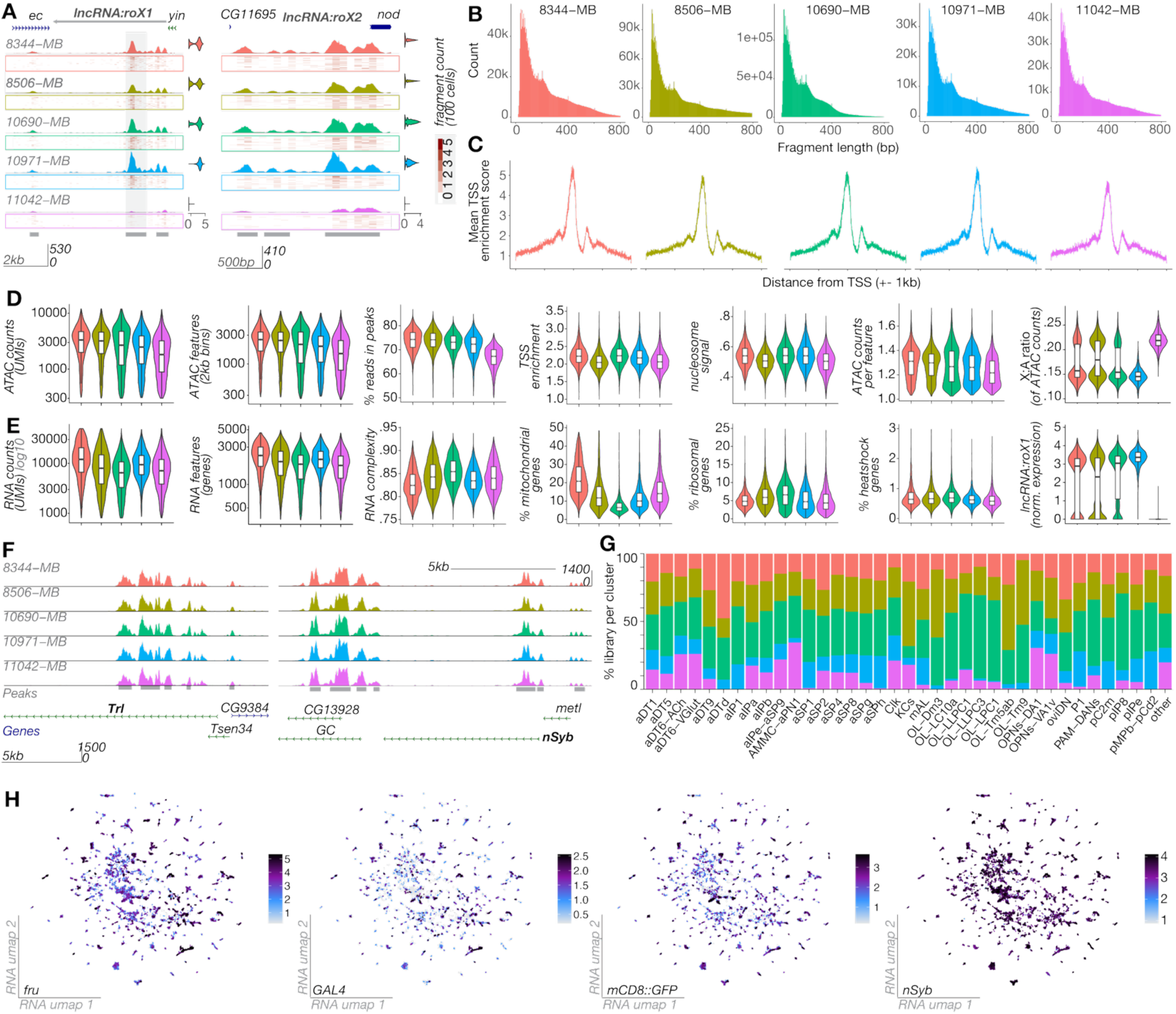
A) Coverage tracks of *lncRNA:roX1*and *lncRNA:roX2* loci per independent replicate. Gray bars represent consensus peaks. B) ATAC fragment length plots per replicate. C) TSS enrichment scores per replicate. D-E) Violin plots of ATAC (D) and RNA (E) QC metrics of filtered cells. Y-axis of counts and features is in log10-scale. F) Coverage tracks of neuron-specific loci *Trl* and *nSyb* split by library replicates. G) Percent of library contribution to annotated clusters. H) Benchmark transcript expression across clusters.

**Supplemental Figure 4, related to Figure 3.**
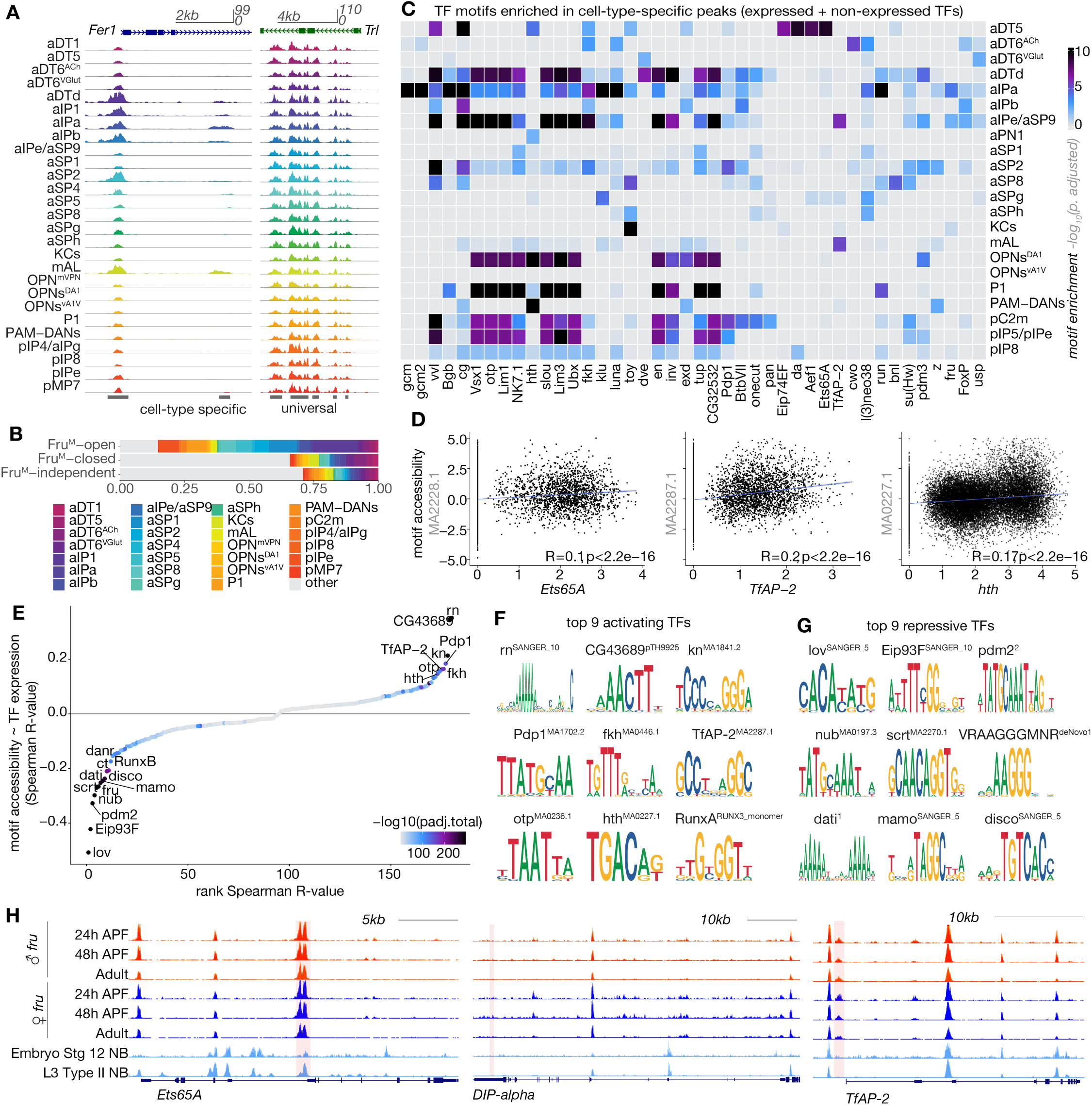
A) Cell-type pseudobulk coverage tracks of cell-type specific transcription factor *Fer1* and ubiquitously expressed *Trl/*GAGA factor. B) Proportion of peaks assigned to annotated clusters in Figure 3C. C) Full heatmap of Figure 3D to include motif enrichment of TFs not expressed in the particular cell types. D) Correlation of TF expression with chromVAR motif accessibility scores per cell to show TF mode-of-action for activating transcription factors. Spearman R-values and correlation p-values are listed. E) Spearman R-values against rank of R-value of expressed transcription factors with their motifs, filtered by dose ∼ expression p-value < 0.01. F) PWMs of top 9 positive TF ∼ motif open chromatin correlations. G) PWMs of top 9 negative TF ∼ motif open chromatin correlations. H) UCSC genome browser screenshots of bulk ATAC coverage from sorted *fru* neurons at 24h, 48h, and adult stages with neuroblast-specific bulk ATAC coverage to show timing of accessibility of genes in Figure 3 G-H.

**Supplemental Figure 5, related to Figure 4.**
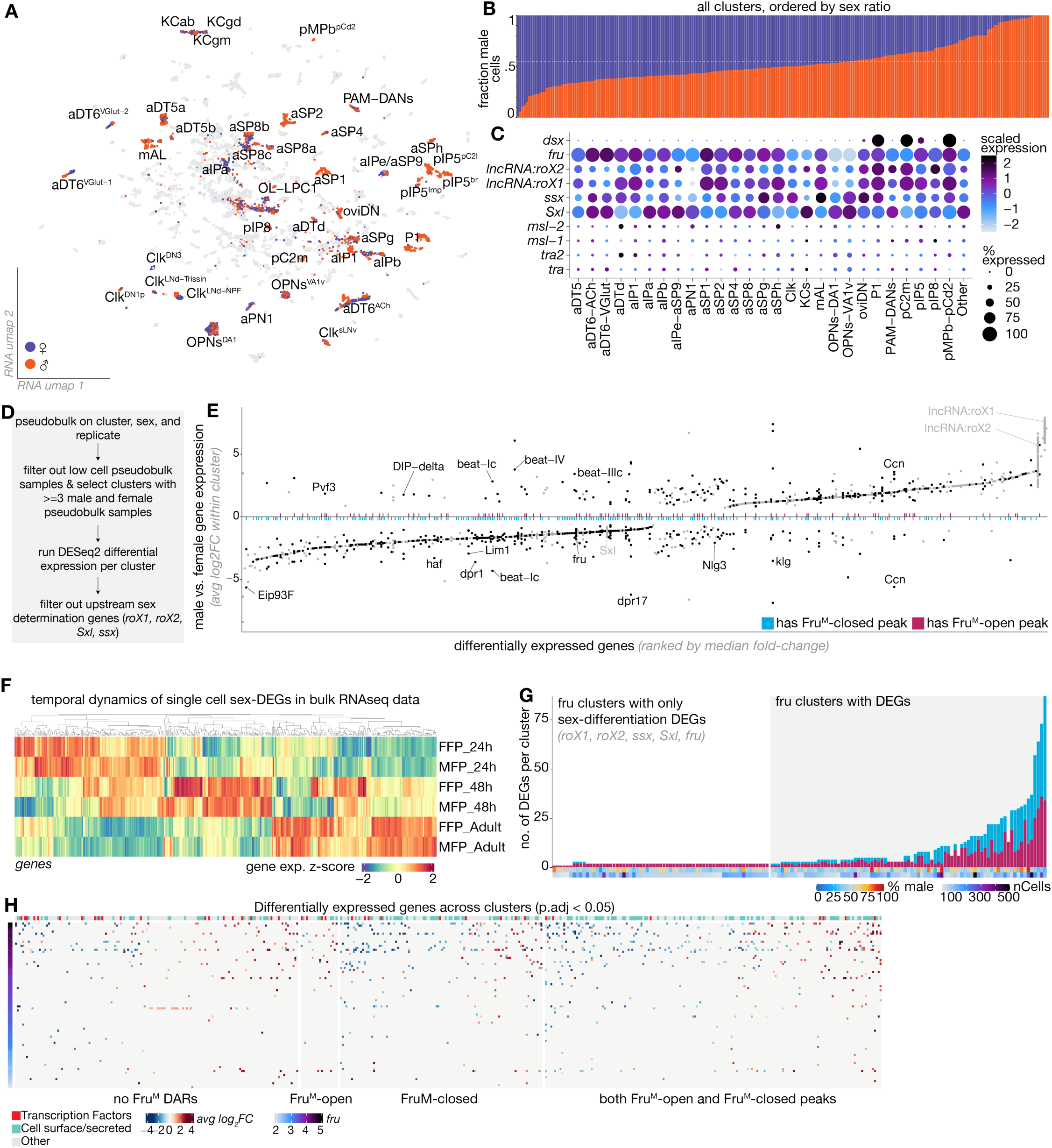
A) RNA umap projection of the full multiome dataset, with sex colored for annotated *fru* neuron subset. B) Fraction of male cells across all clusters. C) Expression of sex-differentiation genes across annotated cell types. D) Flowchart of methodology for generating pseudobulk replicates for DESeq2 testing in male vs female cells per cluster. E) Plot of all differentially expressed genes across clusters, ordered by median fold-change. Points are colored black if they have a Fru^M^-DAR in that gene, and gray if not. Colored bars indicate if gene has a Fru^M^-closed or a Fru^M^-open peak in bulk ATAC data. Columns with multiple dots (e.g. *DIP-delta*, *klg*) represent differential expression of that gene in multiple cell type clusters. F) Temporal dynamics in bulk RNA-seq of genes with sex-differential expression in scRNAseq. G) Number of differentially expressed genes across all sex-shared clusters, including clusters with only sex differentiation DEGs. H) Heatmap of significantly differential genes, with columns ordered by median fold change per genes and rows ordered by average *fru* expression level per cluster. Columns are split by whether differentially expressed genes contain a Fru^M^ differentially accessible region in the bulk ATAC analysis. Genes that appear in multiple rows are differentially expressed in multiple cell types.

**Supplemental Figure 6, related to Figure 5.**
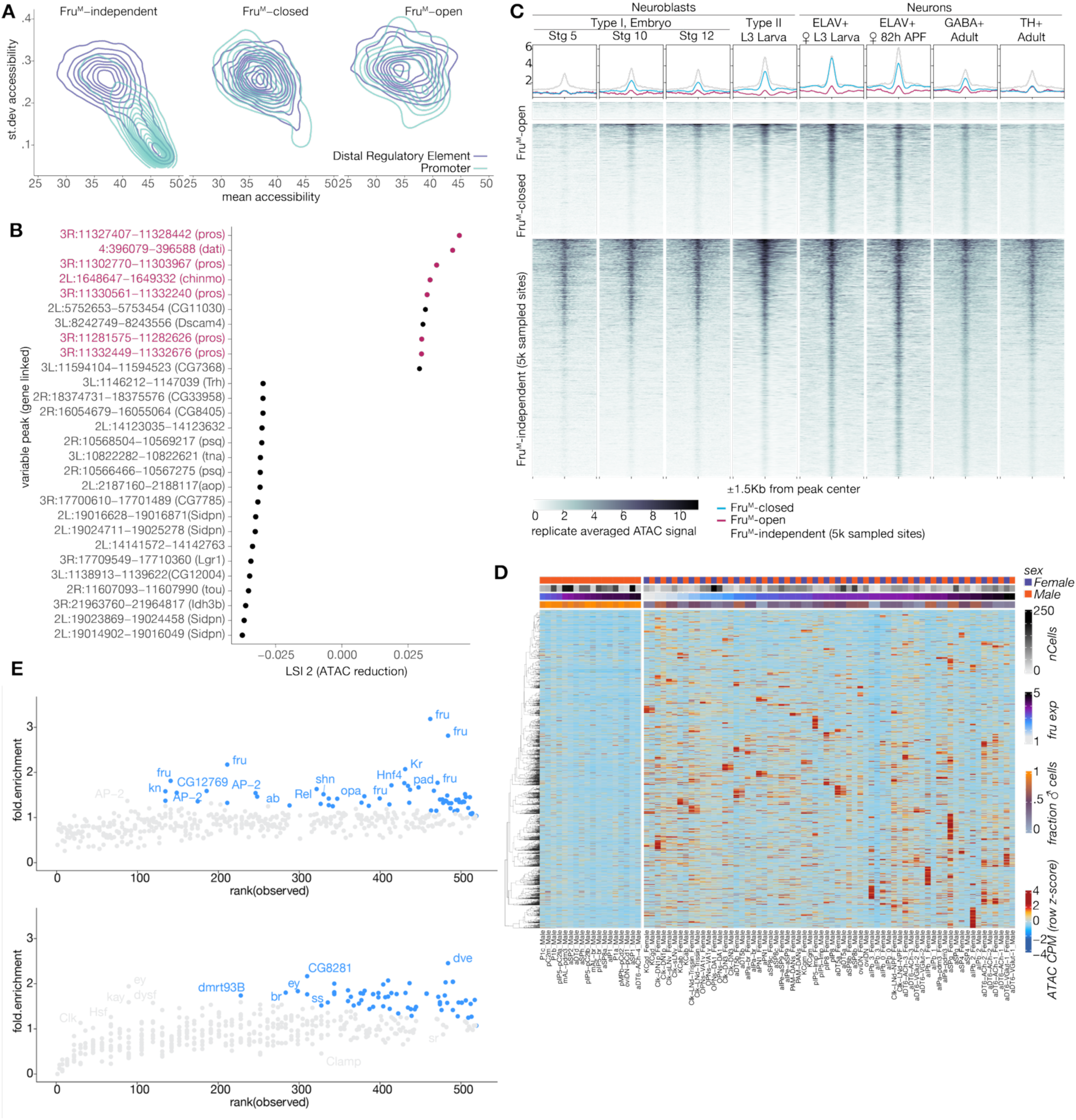
A) Cell-type specificity as standard deviation of region accessibility across 224 clusters against mean accessibility of regions, split by Fru^M^ dependency. B) Top 20 variable peaks contributing to ATAC LSI dimension 2 with nearest gene. C) Signal heatmaps of neuroblast and neuron ATAC-seq signal in Fru^M^-DARs and a 5k random sample of independent peaks. Summary plots show mean accessibility per group. D) Heatmap in Figure 5H with male and female sex-shared cell types interleaved. E) Motif prevalence and enrichment in Fru^M^-closed (top) and Fru^M^-open (bottom) regions using Signac FindMotifs.

**Supplemental Figure 7, related to Figure 6.**
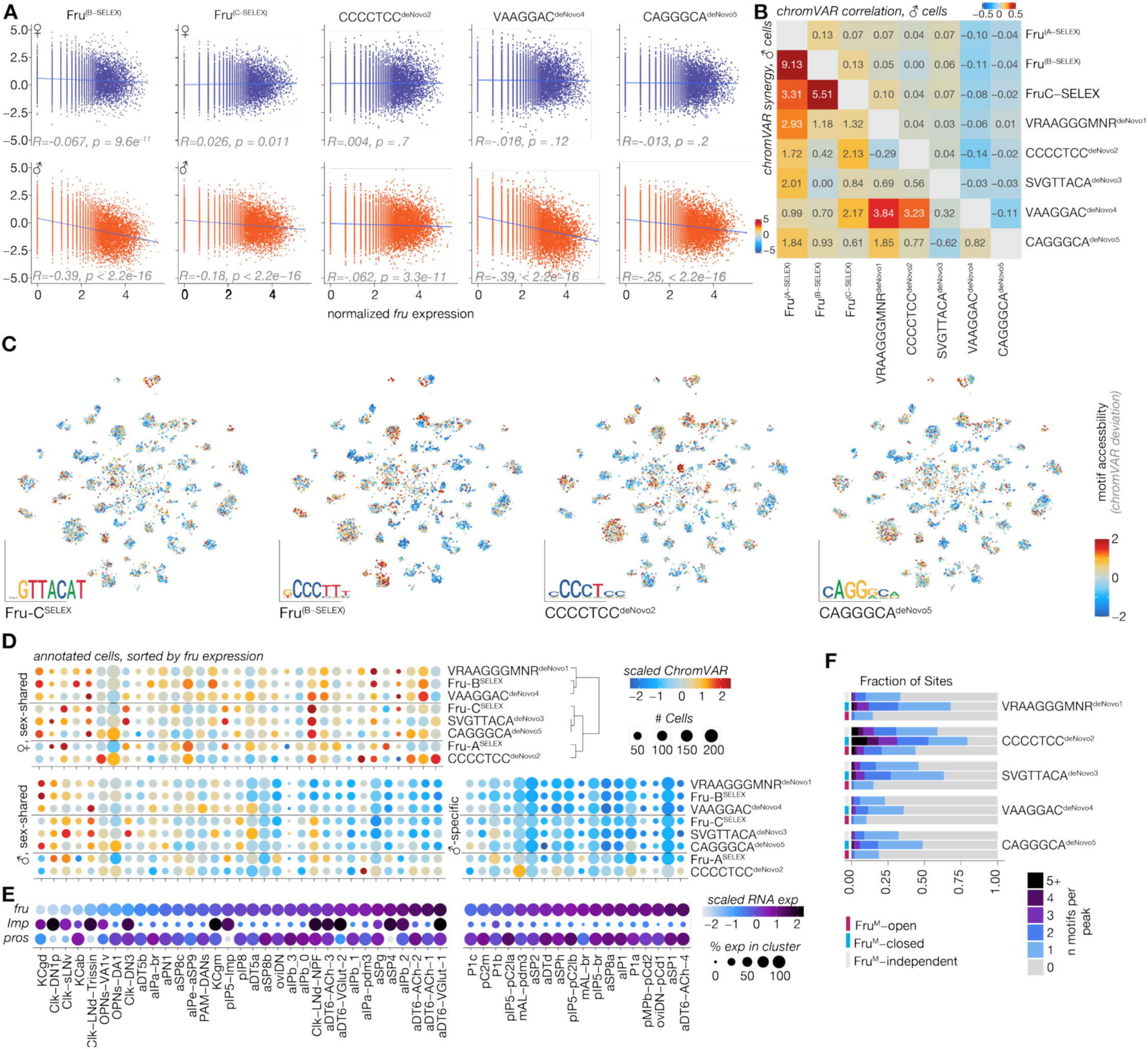
A) Single-cell expression vs chromVAR deviations for additional *de-novo* and SELEX Fru^M^ motifs. B) ChromVAR annotation correlation in male cells representing co-accessibility of motifs and chromVAR annotation synergy in male cells, representing excess cell-cell variability in accessibility for peaks with both motifs present. C) RNA tSNE of chromVAR accessibility scores for additional Fru^M^ motifs across subset annotated cells. D) Dotplot of all Fru^M^ motif chromVAR scores across annotated *fru* cell types. Motifs are ordered by clustering in Fig 6B, clusters are ordered by *fru* expression, with male-specific cell types separated. E) Dotplot of *fru* expression of clusters in D with *Imp* and *pros* expression shown. F) Proportional barplot showing rate of motif multiplicity.

## Notes

### Competing Interest Statement

The authors have declared no competing interest.

